# Insights into vision from interpretation of a neuronal wiring diagram

**DOI:** 10.1101/2023.11.15.567126

**Authors:** H. Sebastian Seung

## Abstract

What insects can see has been probed by over a century of behavioral experiments. Motion and color vision have also been studied through neurophysiology in insect brains. Here I study form vision by interpreting a neuronal wiring diagram of the *Drosophila* optic lobe. The Dm3 “line amacrine” cells are shown to divide into three cell types with oriented dendrites, and to be connected with three TmY cell types, also with oriented dendrites. All six cell types are predicted to respond selectively to oriented visual stimuli, with preferred orientation defined by dendrite orientation. Their receptive fields are predicted by mapping input from other cell types that chiefly convey information from single facets of the compound eye. Dm3 to Dm3 and TmY connectivity is approximated by cross-orientation inhibition and TmY to TmY connectivity by iso-orientation excitation. Both connectivity motifs were previously hypothesized for mammalian visual cortex. Two of the TmY types target a novel type of LC10 cell, which leads by multiple pathways to brain regions that support learning of visual form. Based on the spatial organization of TmY to TmY and LC10 connectivity, I conjecture that flies may see illusory contours and corners.

## Introduction

The question of what insects can see was investigated scientifically starting in the 19th century (Lubbock 1881). For a demonstration that insects can see form (not only color), bees were trained to discriminate between artificial flowers of different shapes (von Frisch 1914). Such behavioral experiments have been refined for over a century, and suggest that insect form vision is primarily for recognizing places rather than objects (Horridge 2009). Starting in the 20th century, insect vision was also investigated by neurophysiologists, who have largely neglected form in favor of motion (Borst and Groschner 2023) and color (Schnaitmann, Pagni, and Reiff 2020) vision. Here I employ a complementary approach that has finally become possible in the 21st century: formulating highly specific predictions and conjectures about vision by interpreting a neuronal wiring diagram.

I begin by defining and analyzing a neural circuit that is intrinsic to the *Drosophila* optic lobe. Half of the circuit is a trio of neuronal cell types, known as Dm3, which are intrinsic to the distal medulla. The other half is a trio of TmY (transmedullary Y) cell types, which project from the medulla to the lobula and lobula plate (Fischbach and Dittrich 1989). The Dm3 trio turns out to have dendrites aligned with three cardinal orientations of the hexagonal lattice of ommatidia in the compound eye. The TmY trio also has dendrites at three distinct orientations.

Visual responses of these cells have not yet been recorded by neurophysiologists. I predict that all six cell types will turn out to be orientation selective, with preferred orientations that correspond with their dendrite orientations. The implication is that the first step of *Drosophila* form vision is a decomposition of the image into small oriented elements like edges or bars, similar to early visual processing in computer science (Marr 1976) as well as models of visual cortex (Fukushima 1980). One further expects that flies perceive orientation rather coarsely, as only three preferred orientations are represented by these neurons.

For more specific predictions, the receptive fields of the six cell types are estimated by mapping their inputs from Tm1 cells. Tm1 cells are just two synapses away from photoreceptors, and Tm1 receptive fields are roughly the width of a single ommatidium. The typical Dm3 receptive field is predicted to be a 3×1 configuration of ommatidia, and the typical TmY receptive field is predicted to be broader.

While Tm1 is the dominant source of their input synapses, Dm3 and TmY cells are also extensively interconnected with each other. Dm3-Dm3 and Dm3-TmY connectivity are interpreted as cross-orientation inhibition, a connectivity motif that has been hypothesized to sharpen orientation tuning in primary visual cortex. Visual stimulation outside the classical receptive field (CRF) is expected to suppress responses to visual stimulation inside the CRF, in a manner that can be predicted from the spatial organization of the inhibition.

TmY-TmY connectivity is interpreted as iso-orientation excitation, another connectivity motif that has long been hypothesized for primary visual cortex. Such excitatory connectivity should at minimum lead to “beyond the CRF” effects, but might even be strong enough to alter the CRF itself. Lateral excitation between parallel TmY neurons could enhance or create position invariance, similar to previous theories of complex cells in primary visual cortex (Chance, Nelson, and Abbott 1999). Furthermore, lateral excitation between collinear TmY neurons could support robust continuation of noisy contours or completion of illusory contours, similar to reports in monkey striate cortex (V1) (Grosof, Shapley, and Hawken 1993) and extrastriate cortex (V2) (von der Heydt, Peterhans, and Baumgartner 1984).

Two of the TmY types synapse onto a novel LC10e cell type, which is conjectured to detect a conjunction of the preferred stimuli of the two TmY types. Assuming this is the case, LC10e could function to detect corners or other kinds of junctions. LC10e is contrasted with the LC15 neuron, which receives input from all three TmY types, and was previously shown to be activated by bars of any orientation (Städele et al. 2020).

Like all LC10 types, LC10e projects to the anterior optic tubercle (AOTu), which leads by multiple pathways to the central complex, a region that is necessary for visual place learning (Wang et al. 2008; Pan et al. 2009; Ofstad, Zuker, and Reiser 2011). LC10e also synapses onto VES044 cells (Scheffer et al. 2020; Schlegel et al. 2023), which project from the lobula to the Vest (VES) neuropil, adjacent to the central complex (Wolff and Rubin 2018).

According to the above predictions, Dm3 and TmY have OFF receptive fields, since their strongest input Tm1 is an OFF cell. It turns out, however, that Dm3 and TmY also receive weaker ON-OFF inputs.

The ON-OFF input regions are similar to the predicted OFF receptive fields, but spatially broader. The ON-OFF regions might give rise to influences “beyond the CRF,” or might influence the CRF itself.

## Results

### Dm3 receives input from collinear columns

A Dm3 cell looks more like a snake than a tree. Its dendrite runs tangentially in layer 3 of the distal medulla making only small side branches. The straightness of the dendrite is the origin of the name “line amacrine cell” (Fischbach and Dittrich 1989; Strausfeld 1970). Later on, Dm3 was divided into two types (Nern, Pfeiffer, and Rubin 2015), which were studied using transcriptomics (Özel et al. 2021; Kurmangaliyev et al. 2020).

It turns out that Dm3 actually comes in three types, with dendrites pointing in three directions (Matsliah et al. 2023). Dm3p dendrites are posteroventral (Fig. 1a) and Dm3q dendrites are posterodorsal (Fig. 1b), as reported previously (Nern, Pfeiffer, and Rubin 2015). The novel type is Dm3v, which has ventrally directed dendrites (Fig. 1c). Dm3p and Dm3q each consist of roughly 450 cells. Dm3v is less numerous, with about 300 cells. These and all subsequent cell counts are for the right optic lobe of a single female adult fly (Methods) (Matsliah et al. 2023).

**Figure 1.**
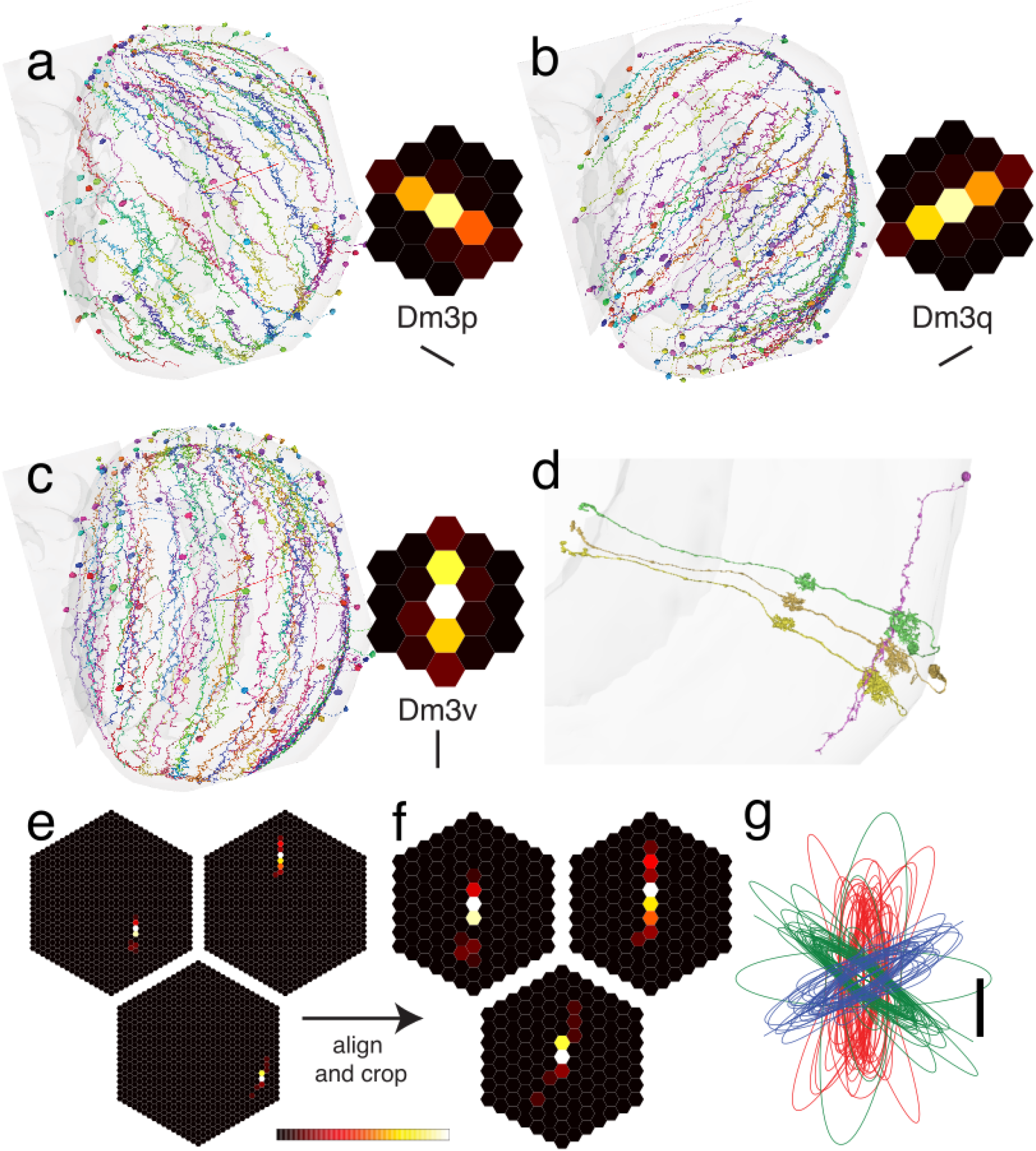
Dm3 cell types and their Tm1 input maps. (a) Dm3p dendrites point in the posteroventral direction. (b) Dm3q dendrites point in the posterodorsal direction. (c) Dm3v dendrites point in the ventral direction. (d) Dm3v cell (pink, 720575940609318851) with three presynaptic Tm1 cells. All presynaptic cells making at least five synapses onto the Dm3v cell are shown. (e) Map of presynaptic Tm1 cells for three example Dm3v cells. Hexel color represents the number of synapses made by the Tm1 cell at that location onto the Dm3v cell. (f) Same Tm1-Dm3v maps as in (e) after aligning on the maximum hexel and cropping. The aligned maps for all Dm3v cells were averaged to create the inset in (c). Average Tm1-Dm3p and Tm1-Dm3q maps were computed similarly, and are the insets in (a) and (b). (g) Tm1 input maps for Dm3v (red), Dm3p (green), and Dm3q (blue) summarized as ellipsoids with parameters from PCA. For the average Tm1-Dm3 maps in (a), (b), and (c), the maximum value in the colormap (white) is 10.6 synapses. Scale bar in (g) is spacing between neighboring columns.

The strongest input to Dm3 is Tm1 (Fig. S1). Dm3 is in turn one of the strongest outputs of Tm1, outranked only by three Pm types (Fig. S2d). Fig. 1d shows a typical Dm3 cell receiving strong Tm1 input in three collinear columns (Fig. 1d), even though the dendrite is long enough to extend over many columns (Fig. 1a-c). For a systematic description, the Tm1 cells presynaptic to each Dm3v cell were mapped (Fig. 1e). The hexels (hexagonal pixels) in the lattices represent medulla columns or ommatidia. The color of each hexel indicates the number of synapses received by the Dm3v cell from Tm1 cells. Aligning (Fig. 1f) and averaging all maps yields the average Tm1-Dm3v input map shown in the inset of Fig. 1a.

This procedure was repeated for all three Dm3 types. The resulting average maps (insets Figs. 1a-c) show that the three Dm3 types are aligned to the three cardinal orientations of the hexagonal lattice. One of the orientations is vertical (*v*). The other two orientations (*p* and *q*) flank the vertical orientation (Zhao et al. 2022; Dickson, Straw, and Dickinson 2008). As observed by (Zhao et al. 2022), the *p* and *q* axes are close to orthogonal in the medulla, which is squashed along the anterior-posterior direction.

The *p* and *q* axes are closer to 120 degrees apart in the eye, where the ommatidia approximate a regular hexagonal lattice.

The hexel images in the figures are drawn on a regular hexagonal lattice, so they should be stretched in the vertical direction when modeling the medulla. They should be left-right reflected when modeling the eye, because the first (outer) optic chiasm connects anterior lamina with posterior medulla, and vice versa (Fischbach and Dittrich 1989). In other words, the *p* and *q* axes are swapped in the eye relative to the medulla.

Each average map is concentrated in a 3×1 array of ommatidia (insets Figs. 1a-c). Some variability, however, is seen in the example maps for single Dm3v cells shown in Fig. 1e. For visualization of the variability across cells, each Tm1-Dm3 connectivity map was treated as a probability distribution and principal components analysis was applied in 2D. The results are graphed as ellipses (Fig. 1g).

### Dm3 receptive field estimates

Physiological recordings of Dm3 cells have not been reported, but Tm1 visual responses have been recorded by a number of labs (Strother, Nern, and Reiser 2014; Behnia et al. 2014; Yang et al. 2016). The Tm1 receptive field is radially symmetric, with a center that is about one ommatidium wide (Arenz et al. 2017). Because this is so narrow, the Tm1-Dm3 connectivity maps (Figs. 1a-c) can be regarded as estimates of Dm3 receptive fields, assuming that a Dm3 cell sums inputs from its presynaptic Tm1 cells. Since the receptive fields are oriented, it follows that Dm3 cells prefer stimuli at the three cardinal orientations (*v*, *p*, and *q*).

Although Tm1 is the strongest input to Dm3 by a large margin (Fig. S1a), Dm3 also receives substantial input from some other numerous (>700 cells) types that could also contribute to the Dm3 receptive field. Maps of their connectivity with Dm3 look similar to Tm1-Dm3 maps (Fig. S3). Like Tm1, most of these inputs (Mi4, Tm2, L3, and Tm9) are known to have receptive fields that are roughly a single ommatidium in width (Serbe et al. 2016; Arenz et al. 2017; Drews et al. 2020). Therefore these inputs are not expected to change the shape of the receptive field estimated from Tm1.

I should clarify that these predictions are for the “classical receptive field” (CRF), defined as that area of the visual field or the retina where visual stimulation can activate the cell (Allman, Miezin, and McGuinness 1985). Classically this was a flashed or moving spot or bar, either light or dark. For some visual neurons, stimulation of areas outside the CRF may modulate the response to stimulation of the CRF (Allman, Miezin, and McGuinness 1985). Such “beyond the CRF” effects will be predicted later based on connectivity between Dm3 cells. For now I will assume that Dm3 input to Dm3, though fairly strong (Fig. S1a), does not contribute to the CRF. It seems safe to assume this because inhibitory input can only suppress activity, and there is no activity to suppress if the CRF is not receiving stimulation.

Based on physiology (Arenz et al. 2017) and predicted neurotransmitters, Tm1 is transient OFF excitatory, Mi4 is sustained ON inhibitory, Tm2 is transient OFF excitatory, L3 is sustained OFF excitatory (Drews et al. 2020), and Tm9 is sustained OFF excitatory. These inputs are consistent with the prediction that Dm3 is an OFF cell.

### Tm1 input to TmY cells

While Dm3 is intrinsic to the distal medulla, TmY cells extend over multiple neuropils, arborizing in the medulla, lobula, and lobula plate (Fischbach and Dittrich 1989). In the distal medulla, TmY4 dendrites have a horizontal orientation, extending symmetrically in two directions on either side of the main trunk (Fig. 2a). There are about 200 TmY4 cells.

**Figure 2.**
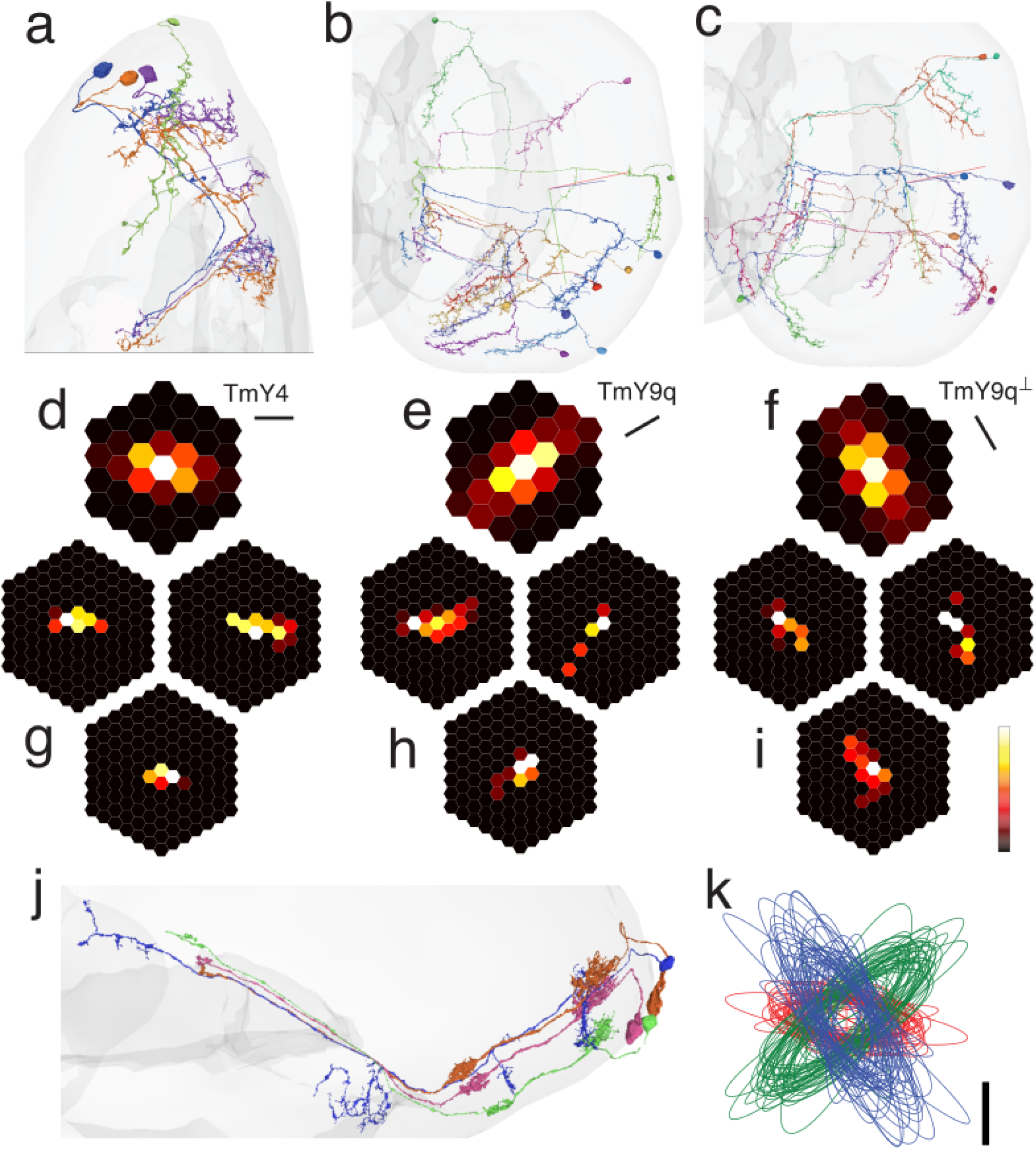
TmY cell types and their Tm1 input maps. (a) TmY4 cells presynaptic to a Dm3v cell (green). TmY4 dendrites in the distal medulla are horizontally oriented and typically extend symmetrically on either side of the main trunk. (b) TmY9q dendrites in the distal medulla point in the anteroventral direction. (c) TmY9q^┴^ dendrites in the distal medulla point in the posteroventral direction. (d) Average Tm1-TmY4 map. (e) Average Tm1-TmY9q map. (f) Average Tm1-TmY9q^┴^ map. (g) Three example Tm1-TmY4 maps, aligned and cropped. (h) Three example Tm1-TmY9q maps, aligned and cropped. (i) Three example Tm1-TmY9q^┴^ maps, aligned and cropped. (j) TmY4 cell (blue, 720575940631972532) with three strong (at least 5 synapses) presynaptic Tm1 partners. (k) Tm1 input maps for TmY4 (red), TmY9q (green), and TmY9q^┴^ (blue), summarized as ellipsoids with parameters from PCA. Maximum value in the colormap (white) is 4.8 synapses for (d), (e), and (f). Scale bar in (k) is spacing between neighboring columns.

TmY9 was originally defined by (Fischbach and Dittrich 1989), and now has been split into TmY9q and TmY9q^┴^ types (Matsliah et al. 2023). In the distal medulla, both TmY9 types have dendrites that are asymmetrically directed to one side of the main trunk. TmY9q dendrites are anteroventrally directed (Fig. 2b), and roughly antiparallel to Dm3q dendrites. TmY9q^┴^ dendrites are posteroventrally directed (Fig. 2c), and orthogonal to Dm3q and TmY9q dendrites. TmY9q^┴^ is bistratified in the lobula, while TmY9q is monostratified (Fig. 2b, c). Variability of TmY9 stratification was previously noted (Tanaka 田中涼介 and Clark 2022), and now can be explained as intertype rather than intratype variation. There are about 180 cells in each of the TmY9q^┴^ ^and TmY9q types.^

Tm1 is one of the top inputs to TmY4 (Fig. S4a, S5a). The average Tm1-TmY4 input map looks horizontally oriented (Fig. 2d), matching the orientation of TmY4 dendrites. The average map is concentrated in a 3×2 array of ommatidia, and extends over a diamond-shaped region with a 5×3 bounding box (Fig. 2d). Input maps for individual TmY4 cells are shown in Fig. 2g. The average input maps for Tm1-TmY9 have similar sizes and aspect ratios, but different orientations (Fig. 2e, f). They extend a bit longer than TmY4 input maps. Example TmY9 cells look similar but exhibit variability (Figs 2h, i). An example TmY4 cell with three strong (at least five synapses) Tm1 presynaptic partners is shown in Fig. 2j. Variation across cells is illustrated by the ellipses drawn in Fig. 2k.

The maps of Tm1 input can be regarded as estimates of receptive fields. TmY4, TmY9q and TmY9q^┴^ are predicted to prefer visual stimuli at the horizontal, *q*, and *q*^┴^ orientations, respectively.

### Cross-orientation inhibition

Dm3 and TmY cells were introduced separately, but in fact they are extensively interconnected with each other. Dm3 to TmY connectivity seems important, judging from its strength relative to other connections. For each Dm3 type, the strongest output is a corresponding TmY type (Fig. S1b). And for each TmY type, the strongest input is a corresponding Dm3 type (Fig. S4a, c).

The correspondences are evident in a visualization of the total number of synapses from every Dm3 type to every TmY type (Fig. 3a). It turns out that Dm3 to TmY connectivity is almost always between dendrites with different orientations. For example, TmY4 has horizontal dendrites, and receives input from Dm3v, which has vertical dendrites. Similarly, TmY9q prefers input from Dm3p, and TmY9q^┴^ prefers input from Dm3q (Fig. 3a).

**Figure 3.**
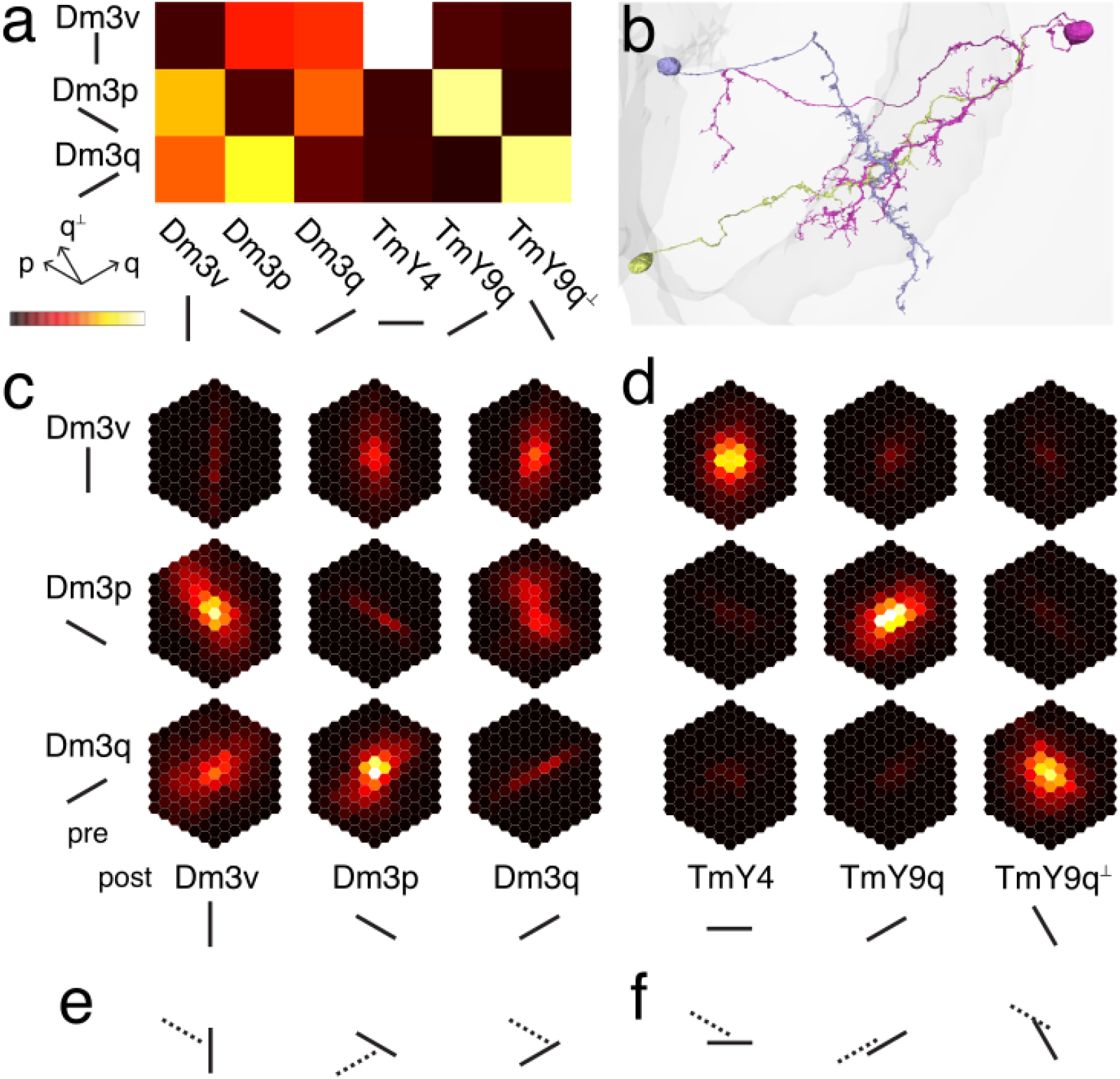
Dm3-Dm3 and Dm3-TmY inhibition. (a) Total number of synapses from Dm3 types (rows) to Dm3 and TmY types (columns). Next to each type name is a line segment indicating the orientation of its dendrites in the distal medulla. Dm3 cells avoid making synapses onto Dm3 cells of the same type (dark diagonal in left 3×3 block). Dm3 cells prefer making synapses onto TmY cells with orthogonal or roughly orthogonal dendrites (bright diagonal in right 3×3 block). (b) Dm3p cell (blue) with presynaptic Dm3q (yellow) and postsynaptic TmY9q (pink). The Dm3q cell disinhibits the TmY9q cell through this pathway, assuming that Dm3 is inhibitory. (c) Average Dm3-Dm3 connectivity maps. Each hexel represents the average number of synapses from a presynaptic cell at that location, assuming the postsynaptic cell is located at the central hexel. The location of each cell is computed from its Tm1 inputs (Methods). (d) Average Dm3-TmY connectivity maps. (e) Cartoon of possible Dm3-Dm3 interactions. For a cell with CRF indicated by the solid line, the dotted line represents a possible beyond-the-CRF region consistent with the spatial organization of connectivity. (f) Cartoon of possible Dm3-TmY interactions. Maximum synapse number in the colormap (white) is 8800 in (a), 1.1 in (c) and 2.9 in (d).

Dm3 to Dm3 connectivity is also important. For each Dm3 type, the second strongest output is another Dm3 type (Fig. S1b). At a population level, Dm3-Dm3 connectivity can be summarized by counting the number of synapses from type to type (Fig. 3a). Each Dm3 type receives fewer synapses from cells of the same type, and more synapses from other Dm3 types (Fig. 3a). In other words, Dm3 cells prefer to synapse onto Dm3 cells of different orientations. This connectivity pattern is known as cross-orientation inhibition.

To summarize the structural analysis, Dm3 output connectivity turns out to be stronger for cells with different dendrite orientations. Fig. 3b illustrates a Dm3 cell synapsing onto another Dm3 cell, which in turn synapses onto a TmY cell. The dendrites of the Dm3 cells are orthogonal. The dendrites of the synaptically coupled Dm3 and TmY cells are also orthogonal.

Dm3 was identified as glutamatergic by (Raghu and Borst 2011), and is predicted by FlyWire (Eckstein et al. 2020) to be either glutamatergic or GABAergic. These neurotransmitters are typically inhibitory in the fly brain, so Dm3 will be assumed inhibitory. Therefore Dm3 output connectivity conforms to a motif known as cross-orientation inhibition, which was introduced by studies of visual cortex dating back 50 years (Benevento, Creutzfeldt, and Kuhnt 1972; Morrone, Burr, and Maffei 1982). Cross-orientation inhibition is well-studied in neural network theory (Ben-Yishai, Bar-Or, and Sompolinsky 1995), and can have the effect of sharpening orientation tuning. If Dm3 types with different preferred orientations inhibit each other, that could have the effect of silencing Dm3 responses to nonpreferred stimuli.

The effects of spatial separation are characterized in the 3×3 array of average Dm3-Dm3 connectivity maps shown in Fig. 3c. Each of these was computed much like the average Tm1-Dm3 and Tm1-TmY connectivity maps of Figs. 1 and 2. Maps for individual cells are given in Data S1. Dm3-Dm3 input maps (Fig. 3c) extend over much larger regions than the CRF estimate for Dm3, suggesting that Dm3-Dm3 connectivity should give rise to “beyond the CRF” effects. For example, according to the Dm3q-Dm3p connectivity map (Fig. 3c bottom middle), stimulating to the upper right or lower left of the CRF of the Dm3p cell is expected to suppress the Dm3p response to stimulation in its CRF, due to the Dm3q inhibitory input to Dm3p. Some beyond-the-CRF effects on each Dm3 type are predicted in Fig. 3e.

Dm3-TmY input maps (Fig. 3d) are somewhat larger than the CRF estimate for TmY, but the difference is not as large, so beyond the CRF effects might be harder to observe. It might be possible to observe disinhibition of TmY cells by Dm3 cells of the same orientation, due to pathways like the one shown in Fig. 3b. Some beyond-the-CRF effects on each TmY type due to disinhibition are predicted in Fig. 3f.

### Iso-orientation excitation

TmY-TmY connectivity is important, judging from the numbers. TmY4/TmY9 is the strongest output of TmY9 cells (Fig. S5d), and the third strongest output of TmY4 cells (Fig. S5b). TmY4/TmY9 is the second strongest input of TmY4 cells (Fig. S5a), and the third strongest input of TmY9 cells (Fig. S5c). At a population level, the organization of TmY-TmY connectivity can be summarized by counting the number of synapses from type to type (Fig. 4a). The strongest TmY-TmY connections are on the diagonal, so that each TmY type prefers to receive synapses from cells of the same TmY type (Fig. 4a).

**Figure 4.**
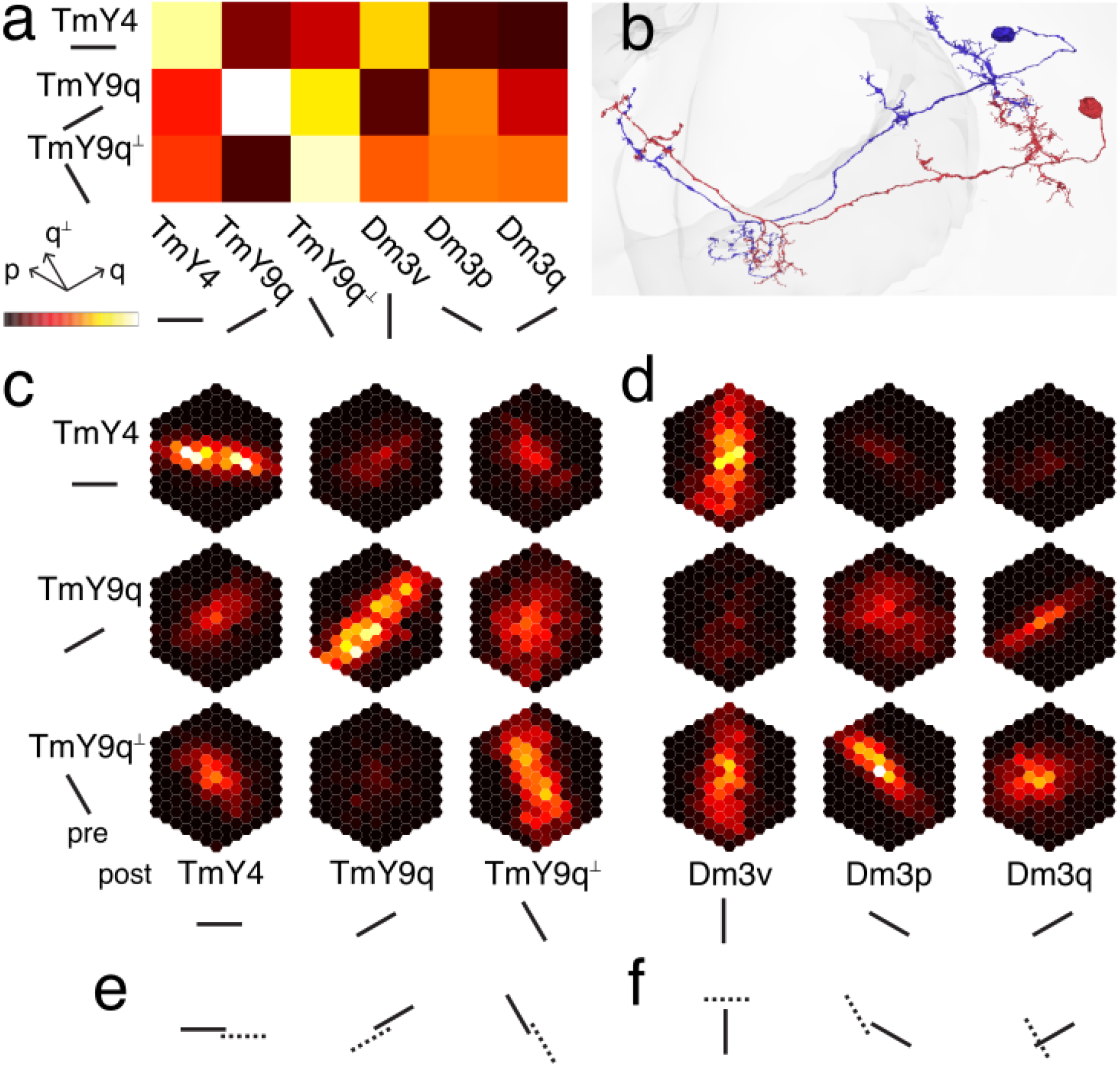
TmY-TmY and TmY-Dm3 excitation. (a) Total number of synapses from TmY types (rows) to TmY and Dm3 types (columns). Next to each type name is a line segment indicating the orientation of its dendrites in the distal medulla. TmY cells prefer to make synapses onto TmY cells of the same type (bright diagonal in left 3×3 block). TmY4 prefers to synapse onto Dm3v, TmY9q prefers to synapse onto Dm3p, but TmY9q^┴^ cells show little preference (right 3×3 block). (b) Pair of TmY4 cells with strong (at least five synapses) and reciprocal connections. (c) Average TmY-TmY connectivity maps. (d) Average TmY-Dm3 connectivity maps. (e) Cartoon of possible TmY-TmY interactions. For a cell with CRF indicated by the solid line, the dotted line represents a possible beyond-the-CRF region consistent with the spatial organization of connectivity. (f) Cartoon of possible TmY-Dm3 interactions. Maximum synapse number in the colormap (white) is 3100 in (a), 0.72 in (c), and 0.29 in (d).

TmY4 is predicted by FlyWire to be cholinergic with high confidence, and will be assumed excitatory. TmY9 has been identified as both cholinergic (Varija Raghu, Reiff, and Borst 2011) and GABAergic (Raghu, Claussen, and Borst 2013). It is predicted by FlyWire to be cholinergic with high confidence, and will be assumed excitatory. With these assumptions, TmY-TmY connectivity conforms approximately to a motif known as iso-orientation excitation.

The dependences of connectivity on spatial separation are characterized in a 3×3 array of TmY-TmY connectivity maps (Fig. 4c) and a 3×3 array of TmY-Dm3 connectivity maps (Fig. 4d). The strongest TmY-TmY connections are on the diagonal, and the maps have orientations that are similar to the orientations of the dendrites. These connections are cartooned in Fig. 4e. The separation vector between the orientation detectors is parallel to the preferred orientations. This spatial configuration is seen in neural network models of contour completion, in which an orientation detector receiving weak or ambiguous input from the image can be driven over threshold by excitation from neighboring orientation detectors.

If the separation vector is orthogonal to the preferred orientations (cartoon not shown), then iso-orientation excitation can lead to more robust orientation tuning (Ben-Yishai, Bar-Or, and Sompolinsky 1995). It could also give rise to positional invariance while preserving orientation selectivity, analogous to complex cortical cells (Chance, Nelson, and Abbott 1999).

Because TmY-TmY connectivity is likely excitatory, it could either lead to beyond-the-CRF effects, or might even be strong enough to modify the CRF itself. On the other hand Dm3 input, because it is inhibitory, was assumed not to modify the CRF.

### Cross-orientation excitation

TmY-Dm3 connectivity is weaker overall than TmY-TmY connectivity (Fig. 4a). Still, Dm3 is one of the strongest outputs of TmY9 (Fig. S5d), and TmY is one of the strongest inputs to Dm3v (Fig. S1c). The organization of TmY-Dm3 connectivity is summarized at a population level in Fig. 4a.

To a first approximation, the connections from TmY to Dm3 mirror those from Dm3 to TmY (see Fig. 3a). In other words, if a Dm3 type inhibits a TmY type, that TmY type reciprocates by exciting the Dm3 type. Since Dm3 to TmY is cross-orientation inhibition, TmY to Dm3 is cross-orientation excitation.

However, this simple approximation is more accurate for TmY4 and TmY9q. TmY9q^┴^ excites all Dm3 types with roughly equal strength. Overall, cross-orientation excitation would be expected to broaden the orientation tuning of Dm3. However, Dm3 likely receives more cross-orientation inhibition than cross-orientation excitation, because TmY4/TmY9 input is substantially weaker than Dm3 input (Fig. S1c).

While there is some reciprocity of excitation and inhibition at the population level, reciprocity does not hold once spatial organization is considered. The shapes of the TmY-Dm3 and Dm3-TmY maps look quite different (Fig. 4d, 3d). Most of the TmY-Dm3 connectivity maps are oriented, while the Dm3-TmY connectivity maps are more isotropic. Some possible spatial configurations of TmY to Dm3 connectivity are cartooned in Fig. 4f.

### LC10e as possible junction detector

LC10 neurons project from the lobula to the anterior optic tubercle (AOTu) of the central brain (Otsuna and Ito 2006). LC10 has been divided into four types (LC10a to LC10d) with distinct stratification profiles in the lobula (Wu et al. 2016; Ribeiro et al. 2018). We identified a novel LC10e type with lobula stratification largely restricted to layer 6. LC10e can be further subdivided into dorsal and ventral variants (Fig. 5a) that have distinct connectivity patterns (Fig. S10).

**Figure 5.**
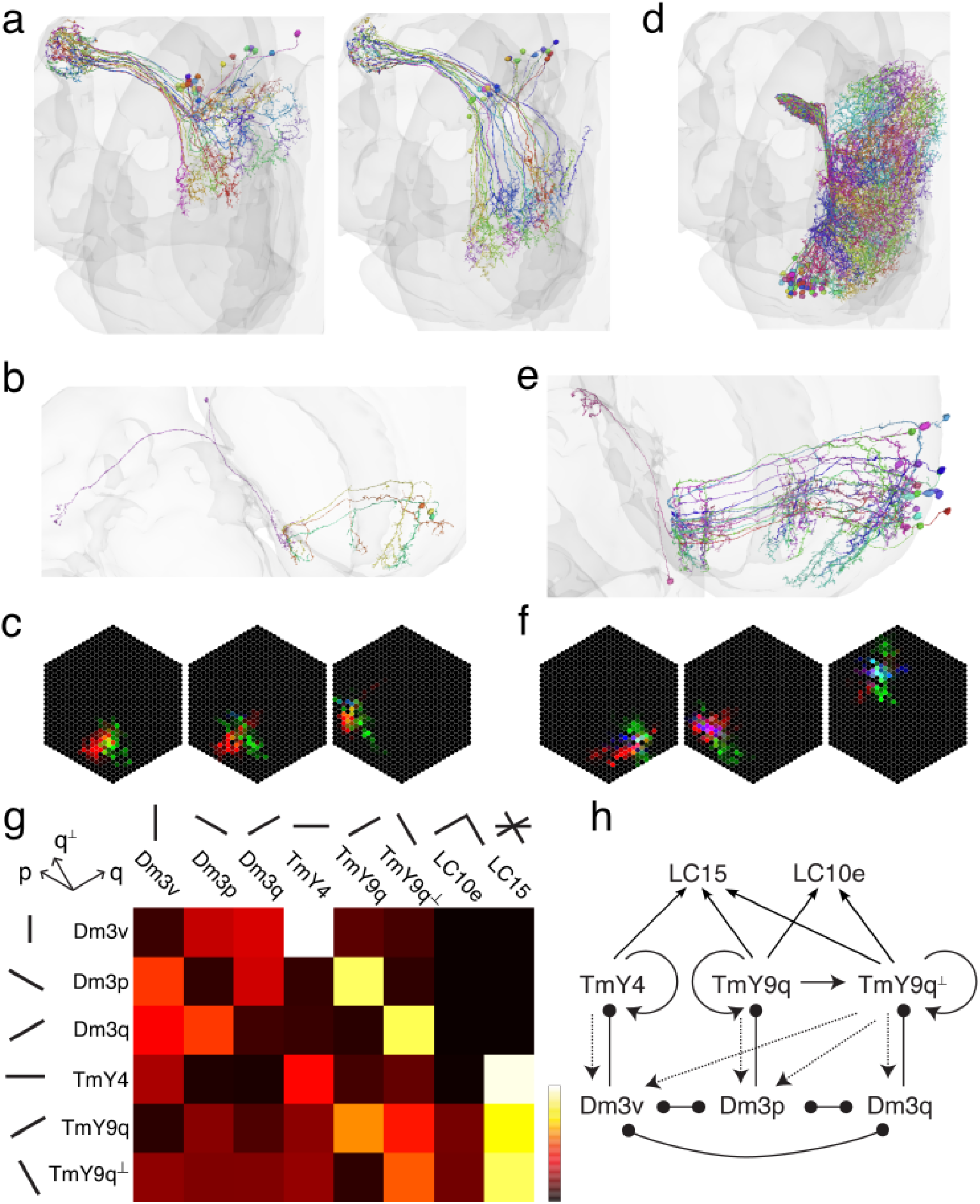
LC cells receiving indirect Tm1 input from TmY cells. (a) LC10e cells project from the dorsal (left panel) and ventral (right panel) lobula to the AOTu (upper left of each panel). (b) An LC10e neuron (pink, 720575940606274592) projecting from the ventral lobula (middle) to the AOTu (far left). Also shown are strong (at least five synapses) presynaptic TmY9q and TmY9q^┴^ partner neurons. Their dendrites in the distal medulla (right) receive Tm1 inputs. (c) Example Tm1-TmY-LC10 maps showing the locations of Tm1 cells providing input mediated by TmY9q (red), TmY9q^┴^ (green), and TmY4 (blue) cells. The red value of each hexel is given by multiplying the number of synapses from Tm1 to TmY9q by the number of synapses from TmY9q to LC10, and the green and blue values are computed similarly. The first two examples resemble a corner and a T-junction. The third example has little form. (d) LC15 cells project from the lobula to PVLP (upper left corner of colored region). (e) An LC15 neuron (pink, 720575940604229152) projecting from the lobula (middle) to the PVLP (upper left). Also shown are strong (at least five synapses) presynaptic TmY4, TmY9q, and TmY9q^┴^ partner neurons. Their dendrites in the distal medulla (right) receive Tm1 inputs. (f) Example Tm1-TmY-LC15 maps. (g) Number of synapses between Dm3, TmY, and LC populations, normalized by geometric mean of presynaptic and postsynaptic population sizes. (h) Cartoon of connectivity in (g). Arrows are excitatory and circles are inhibitory. For each map in (b) and (d), the saturation level of the RGB channels was set at the 95th percentile of nonzero values. The saturation level was 48, 31, and 30 synapse^2^ in the three maps of (b) and 101, 68, and 81 synapse^2^ in the three maps of (d). Maximum value in the colormap is 33 synapses/neuron in (g).

TmY9q and TmY9q^┴^ synapse onto LC10e (Fig. 5b, S4d, S5d). On average, TmY9 inputs are stronger for ventral than for dorsal LC10e (Fig. S10). For clues about the visual features detected by LC10e neurons, it is helpful to map Tm1 cells that are presynaptic to TmY cells that are presynaptic to an LC10e cell. The hexels are color coded depending on whether the intermediary is TmY9q (red), TmY9q^┴^ (green), and TmY4 (blue). The maps for three example ventral LC10e cells are shown in Fig. 5b. There is little or no blue in the maps, since TmY4 makes few or no synapses onto LC10e. The examples resemble corners with the apex pointing upward (Fig. 5c). The maps for dorsal LC10e tend to look noisier, and the shapes are inconsistent (Fig. S6a). The maps for ventral LC10e often resemble corners or T-junctions in the interior of the eye, but are distorted near the edges of the eye (Fig. S6b).

While LC10e receives input from TmY9q and TmY9q^┴^, it is unclear how these two channels interact. One possibility is that LC10e detects a conjunction (logical AND) of activity in the two channels. Based on this assumption, I speculate that ventral LC10e could be a detector of corners or other kinds of junctions, due to the spatial organization in Fig. 5c and S6b.

### Predicted mechanism of LC15 invariance to orientation

LC15 neurons project from the lobula to the posterior ventrolateral protocerebrum (PVLP), a neuropil in the central brain (Fig. 5d) (Wu et al. 2016). All three TmY types synapse onto LC15 (Figs. 5e, S4b, S4d, S5b, S5d). Again it is helpful to map the Tm1 cells that are presynaptic to TmY cells that are presynaptic to an LC15 cell. Maps for example LC15 cells are shown in Fig. 5f, and for all LC15 cells in Fig. S7. Maps generally include all three colors, indicating that all three TmY types synapse onto LC15. For each LC15 cell, the three channels take Tm1 input from eye regions that overlap with each other. No additional spatial organization is evident.

Given that LC15 connectivity to all three orientation channels is indiscriminate, it is natural to hypothesize that LC15 activity is also indiscriminate, detecting a disjunction (logical OR) of activity in the three channels. If TmY orientation tuning is sufficiently broad, it follows that LC15 should be activated by a stimulus of any orientation. Indeed, recordings of LC15 visual responses show that LC15 is activated by bars of any orientation (Städele et al. 2020).

### LC10e pathways to the central complex

Behavioral experiments on form vision in bees have generally involved trained visual discriminations, which require some memory of visual stimuli. The central complex is a brain region necessary for learning visual forms (Wang et al. 2008; Pan et al. 2009; Ofstad, Zuker, and Reiser 2011). There are multiple pathways from AOTu to the central complex. For example, some of the LC10e targets in the AOTu project to the lateral accessory lobe (LAL), a neuropil that provides input to the central complex. Characterization of such pathways is left for future work.

In addition, there is another route for LC10e output to reach the central complex that does not travel through AOTu. A strong output of LC10e is VES044 (Scheffer et al. 2020; Schlegel et al. 2023), which projects from the lobula to the Vest (VES) neuropil (Wolff and Rubin 2018). The LC10e to VES044 synapses are located in the lobula; their arbors do not overlap in the central brain. One of the VES044 cells is dominated by LC10e input, while the other two receive a more even mixture of LC10d and LC10e input.

### ON-OFF input regions

Inputs to Dm3 cells from Tm1, Mi4, Tm2, and L3 were mapped above (Figs. 1, S3). Dm3 has an additional input T2a (Fig. 6a, S1) about which little is known. T2a is predicted to be a transient ON-OFF cell, since its strongest inputs are Tm1 (transient OFF excitatory) and Mi1 (transient ON excitatory) (Fig. S8a).

**Figure 6.**
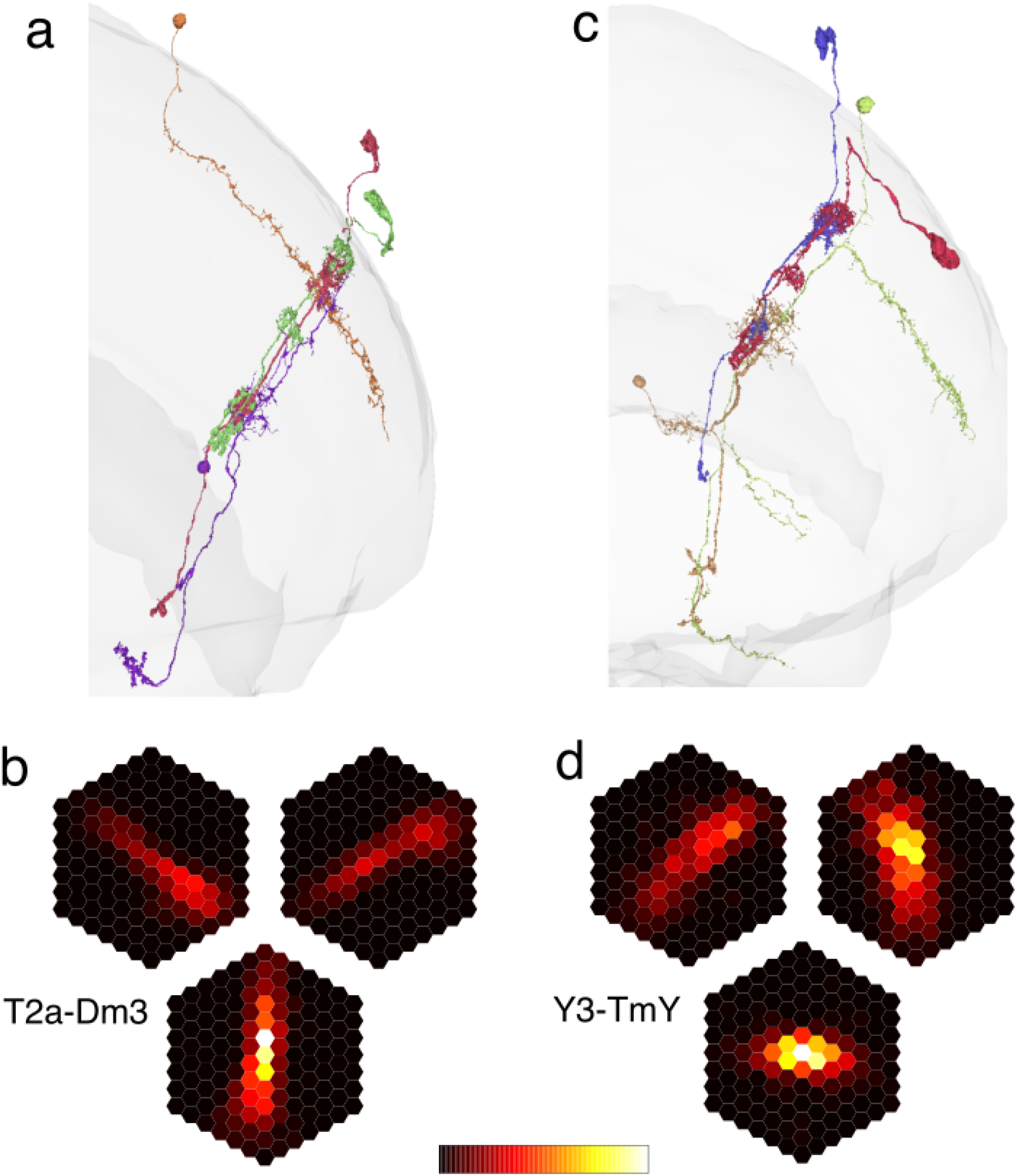
ON-OFF input regions. (a) T2a (purple) with a postsynaptic Dm3v (orange) and presynaptic Tm1 (red) and Mi1 (green). Tm1 is also presynaptic to Dm3v. (b) T2a-Dm3v (bottom), T2a-Dm3p (top left), and T2a-Dm3q (top right) input maps, aligned and averaged. (c) Y3 (orange) with postsynaptic TmY9q (green) and presynaptic Tm1 (purple) and Mi1 (red). (d) Y3-TmY4 (bottom), Y3-TmY9q (top left), and Y3-TmY9q^┴^ (top right), aligned and averaged. Maximum synapse number in the colormap (white) is 0.99 in (b) and 1.5 in (d).

Average T2a-Dm3 input maps are long and thin (Fig. 6b). Tm1-T2a-Dm3 and Mi1-T2a-Dm3 input maps are similar, but broader (data not shown). They define an “ON-OFF input region” of the Dm3 cell, a part of the visual field where both ON and OFF stimuli can influence Dm3 activity. For each Dm3 type, the ON-OFF input region resembles the estimated CRF (Fig. 1a-c), but is longer and thicker.

Y3 (Fig. 6c) is a strong input to TmY4 (Fig. S4a, S5a), and TmY9 cells (Fig. S4b, S5b). Since their original definition by (Fischbach and Dittrich 1989), nothing has been reported about Y3 except that they are targets of T4 and T5 (Shinomiya et al. 2022). Mi1 and Tm1 are the strongest Y3 inputs (Fig. S9a), so Y3 is predicted to be ON-OFF like T2a.

Average Y3-TmY input maps are oriented (Fig. 6f). Tm1-Y3-TmY and Mi1-Y3-TmY input maps are similar, but broader (data not shown). They define an ON-OFF input region for each TmY type, which resembles the estimated CRF (Fig. 2d-f), but is broader. The ON-OFF input regions for TmY9q and TmY9q^┴^ look bent. For TmY9q the dorsal part has the *q* orientation, but the ventral part has the *p*^┴^ orientation (Fig. 6f, top left). For TmY9q^┴^ the dorsal part has the *q*^┴^ orientation, and the ventral part has the vertical orientation (Fig. 6f, top right). This difference in orientation arises because Y3-TmY connectivity in the dorsal part is mainly through synapses in the medulla, while Y3-TmY connectivity in the ventral part is mainly through synapses in the lobula and lobula plate. Evidently the orientation of TmY9 dendrites can vary a little across neuropils. Whether this is merely a quirk of development or has functional significance is unclear.

To summarize, Dm3 and TmY were first predicted to have OFF input regions on the basis of direct Tm1 input. Now they are also predicted to have ON-OFF input regions that are spatially broader, due to indirect Mi1 and Tm1 input. In one scenario, the CRF is given by the OFF input region, and beyond-the-CRF effects arise from the ON-OFF input regions. Alternatively, the ON-OFF input region might even be strong enough to make the CRF larger than the OFF input region.

## Discussion

Dm3 cells were described in *Drosophila* by (Fischbach and Dittrich 1989), and analogous line amacrine cells were described in other insect species well before that (Strausfeld 1970). Such evolutionary conservation contrasts with the pronounced interspecies variability said to be characteristic of most Dm types (Fischbach and Dittrich 1989). Evolutionary conservation of Dm3 is consistent with the idea that orientation selectivity is fundamental to form vision. It also justifies the comparison with bee studies in the Discussion to follow.

The present work also stimulates comparisons with mammals, as cross-orientation inhibition was hypothesized for primary visual cortex over 50 years ago (Benevento, Creutzfeldt, and Kuhnt 1972; Morrone, Burr, and Maffei 1982). Most empirical tests have utilized voltage clamp recordings to disentangle excitatory and inhibitory input, and the results are still debated (Isaacson and Scanziani 2011). Iso-orientation excitation was also proposed long ago for primary visual cortex (Mitchison and Crick 1982). Such connectivity by definition exists for neurons in the same orientation column, but for more distant neurons the experimental evidence remains equivocal in spite of decades of studies (Chavane, Perrinet, and Rankin 2022).

Cross-orientation inhibition (Dm3 to Dm3 and Dm3 to TmY) and iso-orientation excitation (TmY to TmY) are now empirical facts in the fly optic lobe (Figs. 5g, h), if “orientation” refers to dendritic structure. They remain predictions for now if the connectivity motifs refer to preferred orientation, since the visual responses of the neurons are still unmeasured. The weaker TmY to Dm3 connectivity (Fig. 5g) is approximately described by cross-orientation excitation, which was not previously proposed for cortex.

The data can also be used to quantify deviations from the idealized descriptions. For example, the off-diagonal TmY to TmY connections deviate from iso-orientation excitation. And TmY9q^┴^ synapses roughly equally on all Dm3 types, deviating from cross-orientation excitation. Importantly, the data allows precise quantification of the spatial organization of cross-orientation inhibition (Figs. 3c, d) and iso-orientation excitation (Figs. 4c, d). Such spatial organization is crucial for understanding the implications of these connectivity motifs for visual function. In principle, similarly thorough and conclusive analyses should be possible with the MICrONS cortical dataset that combines calcium imaging and EM reconstruction, but the number of fully proofread neurons in this dataset is relatively small at present (MICrONS Consortium et al. 2021).

### Motion-form interactions

Behavioral experiments with honeybees were used to argue that the orientation of a visual stimulus is computed independently from its direction of motion (Mandyam V. Srinivasan, Zhang, and Rolfe 1993). Indeed, Dm3 inputs and outputs do not contain any cell types known to be involved in motion computation (Fig. S1). However, TmY4 receives input from T4c, T5c, T4d, and T5d (Figs. S4, S5).

These neurons are activated by upward and downward motion, and are also known to prefer horizontally oriented stimuli (Fisher, Silies, and Clandinin 2015). Therefore, input from the motion system is expected to enhance the OS of TmY4, and also cause TmY4 responses to be stronger for moving stimuli than for flashed stimuli. Since T4 and T5 are ON and OFF, they define an ON-OFF input region to TmY4. This is potentially stronger than the T2a-mediated ON-OFF input region to TmY4, judging from synapse numbers (Fig. S4, S5). TmY9q and TmY9q^┴^ receive input from T5a and T4a (Figs. S4, S5), but this pathway is rather weak.

In addition to the influence of motion on form computation, there also exist pathways for the orientation selective circuit to influence motion computation, as TmY4 synapses onto LPi⇐T5c⇐T5d cells (Figs. S4, S5), which locally inhibit T4c, T5c, T4d, and T5d. TmY9q and TmY9q^┴^ synapse onto a number of types (Y1, Y11, Y12, etc.) that are likely to be DS because they are directly downstream from T4 and T5 cells.

TmY4 receives inhibition from several lobula plate intrinsic types, including the full-field cells LPi⇐T5a⇒H2 and LPi⇐T5b⇒DCH, which are predicted to be GABAergic/inhibitory. This suggests that TmY4 might be suppressed by horizontal background motion. TmY9q^┴^ (and to a lesser extent TmY9q) receives inhibition from LPi⇐T5c⇐T5d, suggesting that TmY9q^┴^ (and TmY9q) might be suppressed by vertical background motion.

### OFF input regions as receptive field predictions

OFF input regions for Dm3 and TmY cells were interpreted as receptive field estimates, which can be viewed as predictions to be tested by future visual physiology experiments. Alternatively, the receptive field estimates can be viewed as confirmations of 30-year-old predictions based on bee behavior that there should be “three broadly tuned orientation-sensitive channels” (Mandyam V. Srinivasan, Zhang, and Rolfe 1993; M. V. Srinivasan, Zhang, and Witney 1994). Orientation selective (OS) neurons were predicted to span a maximum of three ommatidia (Horridge 2003), which matches the OFF input regions of Dm3 cells (Figs. 1, S3), and approximates the OFF input regions for TmY cells (Fig. 2). So the old predictions are confirmed, assuming that the OFF input region is in fact an estimate of the receptive field. Given that Dm3 dendrites are so long, Dm3 receptive fields would not previously have been guessed to extend over just three ommatidia. This prediction depends on mapping the input connectivity of Dm3.

### Interaction between OFF and ON-OFF input regions

Dm3 and TmY are predicted to have ON-OFF input regions that are broader than the OFF input regions (Fig. 6). As mentioned earlier, it is unclear whether the ON-OFF input regions shape the CRF, or whether they only contribute “beyond the CRF” modulatory influences to a CRF determined by the OFF input region.

For TmY types, the OFF input region is determined primarily by synapses received in the medulla, while the ON-OFF input region is determined primarily by synapses received in the lobula and lobula plate. It is unclear how much voltage varies across a TmY cell, but substantial variation may exist in at least one other type of Drosophila central neuron (Gouwens and Wilson 2009). This raises a possible complication that the relative strength of the OFF and ON-OFF input regions might vary across neuropils, and their interaction might involve some kind of nonlinearity.

### Sharpening orientation tuning

Ignoring variability across cells, the OFF input region to Dm3 cells is a 3×1 configuration of ommatidia (Fig. 1, S3). A stimulus at the preferred orientation will activate all 3 ommatidia, while a stimulus at a nonpreferred orientation will activate only a single ommatidium. If this were the only input to Dm3, and Dm3 response were linear, then its nonpreferred response would be one-third the preferred response. There would be even less contrast between nonpreferred and preferred, if one were to take into account that the central ommatidium of the OFF input region has higher weight than the two flanking ommatidia (Fig. 1, S3). The contrast between nonpreferred and preferred responses should be increased by cross-orientation inhibition between Dm3 cells, which would tend to inhibit responses to nonpreferred orientations.

### Position invariance and contour completion

Depending on its strength, iso-orientation excitation between TmY cells could give rise to beyond-the-CRF effects, but could also reshape the CRF. In the latter case, the CRF of a TmY cell would not be given by its Tm1-TmY map, but some combination of the Tm1-TmY maps of coupled TmY cells. For a side-by-side configuration, this would result in position invariance through a mechanism like that hypothesized for complex cortical cells (Chance, Nelson, and Abbott 1999). For a collinear configuration, this could result in TmY responses to illusory contours.

On the other hand, the TmY-TmY interaction might be weak, so that stimulating the Tm1 cells presynaptic to one TmY cell would only be strong enough to enhance the response to stimulating the Tm1 cells presynaptic to the other TmY cell. In this case, the CRF for each TmY cell would be given by its Tm1-TmY map, and the TmY-TmY interaction would result in “beyond the CRF” effects. For a collinear configuration, this would result in enhancing responses to weak or noisy contours, but not responses to completely illusory contours.

Bees have been reported to perceive illusory contours (Horridge, Zhang, and O’Carroll 1992), but the experiment was later declared to be irreproducible (Horridge 2009). The connectivity maps presented here could potentially be used to design illusory contour stimuli, so this topic seems worth reopening.

### Variability across cells of the same type

Average connectivity maps are shown in most figures, and often correspond well with connectivity maps for single cells (Fig. 1e, 2g, 2h, 2i), but there is also variability across cells of the same type. In particular, the connectivity maps for individual LC10e and LC15 cells (Figs. 5b, 5d, S6, S7) exhibit so much variability that averaging them was avoided. Some variability is due to inaccurate 3D reconstruction of cells or detection of synapses. But true biological variability presumably exists too (Takemura et al. 2015). One can imagine that the visual responses of these cells could turn out to be even more variable than the connectivity maps. Alternatively, visual responses might turn out to be robust to variations in connectivity, due to various connectivity motifs mentioned above.

### Feature hierarchies and convolutional nets

Simple and complex cells were hypothesized to be the first level in a feature hierarchy for form vision (Hubel and Wiesel 1962). This idea became the basis of convolutional nets (Fukushima 1980; LeCun et al. 1989), a popular approach to form vision in artificial intelligence. The LC10e type of Fig. 5a is conjectured to detect a conjunction of TmY9q and TmY9q^┴^ activation, specifically visual stimulation of two oriented stimuli arranged in a corner or T-junction (Fig. 5b, S6). The conjecture is specific to the ventral variant of LC10e. This makes sense because the ventral visual field is expected to be more important for form vision, assuming that the fly is above the landmarks or objects to be seen. Whether LC10e activation requires a conjunction of TmY9q and TmY9q^┴^ activation is difficult to predict by analysis of wiring alone. Alternatively, LC10e might detect a disjunction of its inputs. This is likely the case for LC15, and may explain why LC15 detects a bar of any orientation (Städele et al. 2020).

### Neurophysiology and brain simulation

The present work attempted to understand visual function by interpreting a neuronal wiring diagram. Although “line amacrine” cells homologous to Dm3 were observed over half a century ago in *Diptera* (Strausfeld 1970), their visual responses have never been published, and no function has ever been ascribed to them. The visual functions of TmY types are also unknown. Based on anatomy alone, predictions and conjectures were advanced about receptive fields and other visual response properties of Dm3 and TmY neurons.

One might criticize this paper as incomplete, because its predictions are crying out to be tested by neurophysiology. But interpretation of the wiring diagram is arguably a significant contribution in its own right. Attempting to predict receptive field properties before they are measured is unusual and interesting, and a harbinger that it will become standard practice for theorists to use connectomes to provide highly specific predictions that guide physiologists to do worthwhile experiments, rather than merely fitting models after the fact.

One might also complain that this paper is incomplete without modeling the dynamics of neural activity based on the neuronal wiring diagram. Indeed, such a connectome-based approach to brain simulation is being developed by a number of researchers (Lappalainen et al. 2023; Shiu et al. 2023). The present paper shows that interpretation of a wiring diagram can be a useful intermediate step before creating a full-blown model of neural network dynamics. Predictions can already be extracted just from interpretation of the wiring diagram.

### Neural development

As mentioned earlier, Dm3p and Dm3q were previously identified by (Nern, Pfeiffer, and Rubin 2015). They were dubbed Dm3a and Dm3b by (Özel et al. 2021), and shown to have transcriptomes that differ before adulthood (P50 or earlier). Immunostaining showed that Dm3q expresses Bifid, while Dm3p does not. The authors also analyzed a seven column medulla EM dataset (Takemura et al. 2015) and found that Dm3 types prefer to synapse onto each other. This finding foreshadowed the present work, which shows that cross-orientation inhibition includes Dm3v as well as Dm3p and Dm3q. It would be interesting to know how Dm3v differs in gene expression from Dm3p and Dm3q, what molecules guide Dm3 dendrites to grow in three directions, and how Dm3 dendrites “decide” how far to grow, even after turning 90 degrees at the border of the medulla (Fig. 1). The molecules that establish connectivity preferences of Dm3 and TmY cells (Fig. 3a and 4a) are of obvious interest. There should also be molecules that establish preferences for synapse formation at particular dendritic locations. Such preferences are important for the spatial organization of connectivity in Figs. 3c and 4c, but were not analyzed here. In connectivity patterns, Dm3p and Dm3q are more similar to each other than to Dm3v, and TmY9q and TmY9q^┴^ are more similar to each other than to TmY4. These similarities are shown in the dendrogram of Fig. S3 in (Matsliah et al. 2023). It will be interesting to know whether similarity of connectivity corresponds with transcriptomic similarity.

## Supplementary Figures

**Figure S1.**
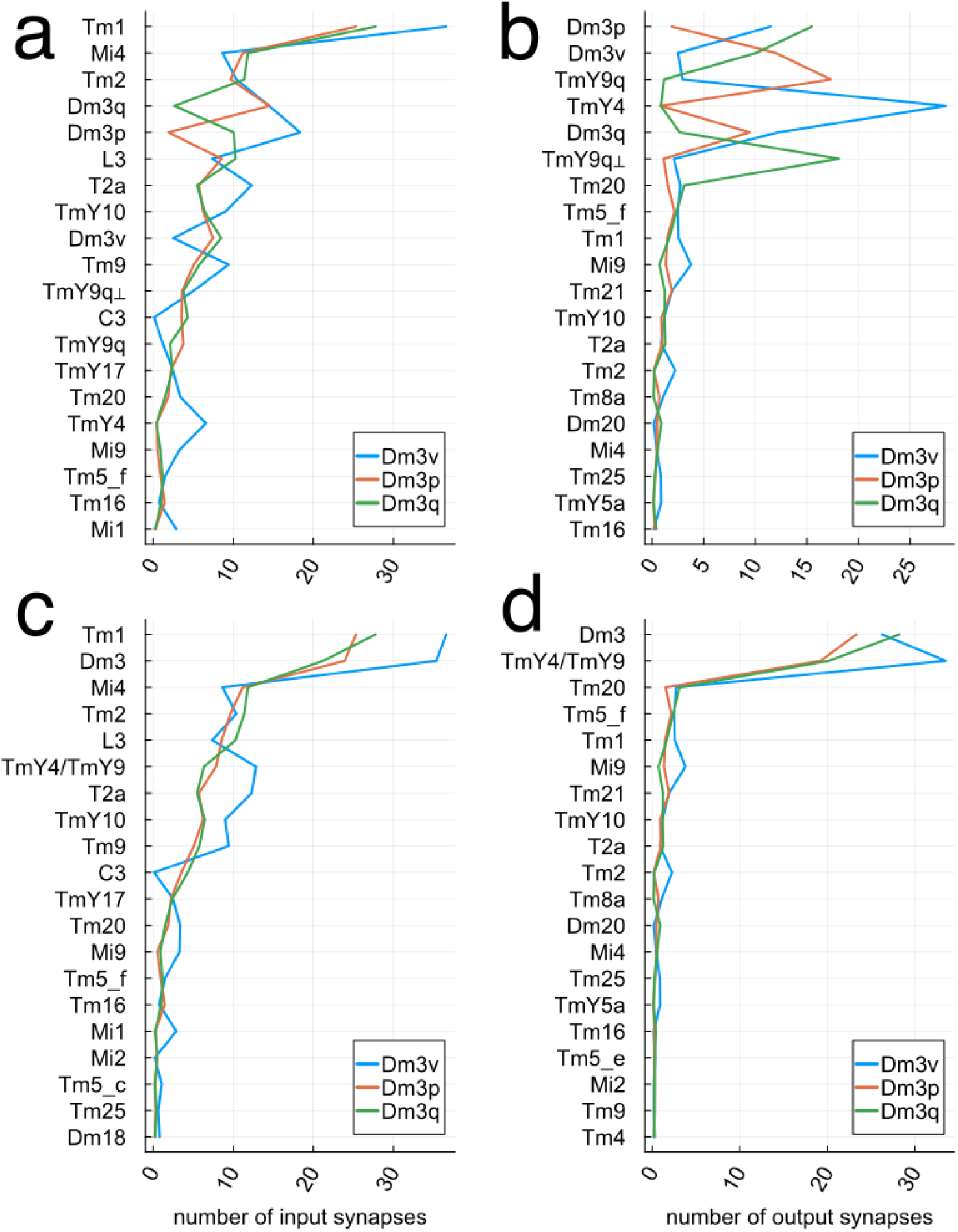
Dm3 inputs and outputs. (a) Average number of input synapses received by a Dm3 cell from presynaptic cell types. Presynaptic types are ordered by the total number of synapses they make onto all three Dm3 types. (b) Average number of output synapses sent by a Dm3 cell to postsynaptic cell types. Postsynaptic types are ordered by the total number of synapses they make onto all three Dm3 types. (c) Same as (a), except Dm3v, Dm3p, and Dm3q are grouped into the presynaptic class Dm3, and TmY4, TmY9q, and TmY9q are grouped into the presynaptic class TmY4/TmY9. (d) Same as (b), except for the groupings Dm3 and TmY4/TmY9.

**Figure S2.**
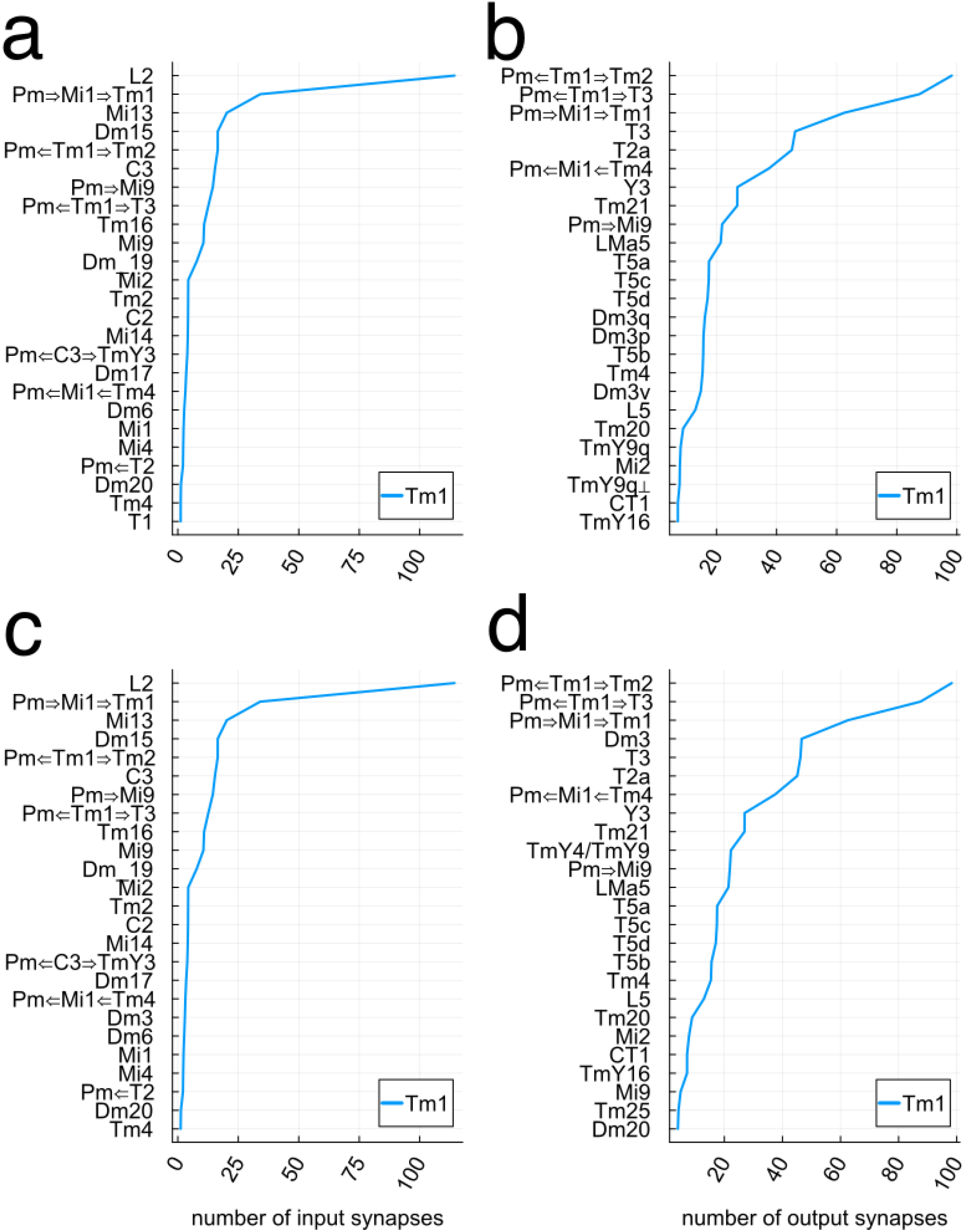
Tm1 inputs and outputs. (a) Average number of input synapses received by a Tm1 cell from presynaptic cell types. (b) Average number of output synapses sent by a Tm1 cell to postsynaptic cell types. (c) Same as (a), except Dm3v, Dm3p, and Dm3q are grouped into the presynaptic class Dm3, and TmY4, TmY9q, and TmY9q are grouped into the presynaptic class TmY4/TmY9. (d) Same as (b), except for the groupings Dm3 and TmY4/TmY9.

**Figure S3.**
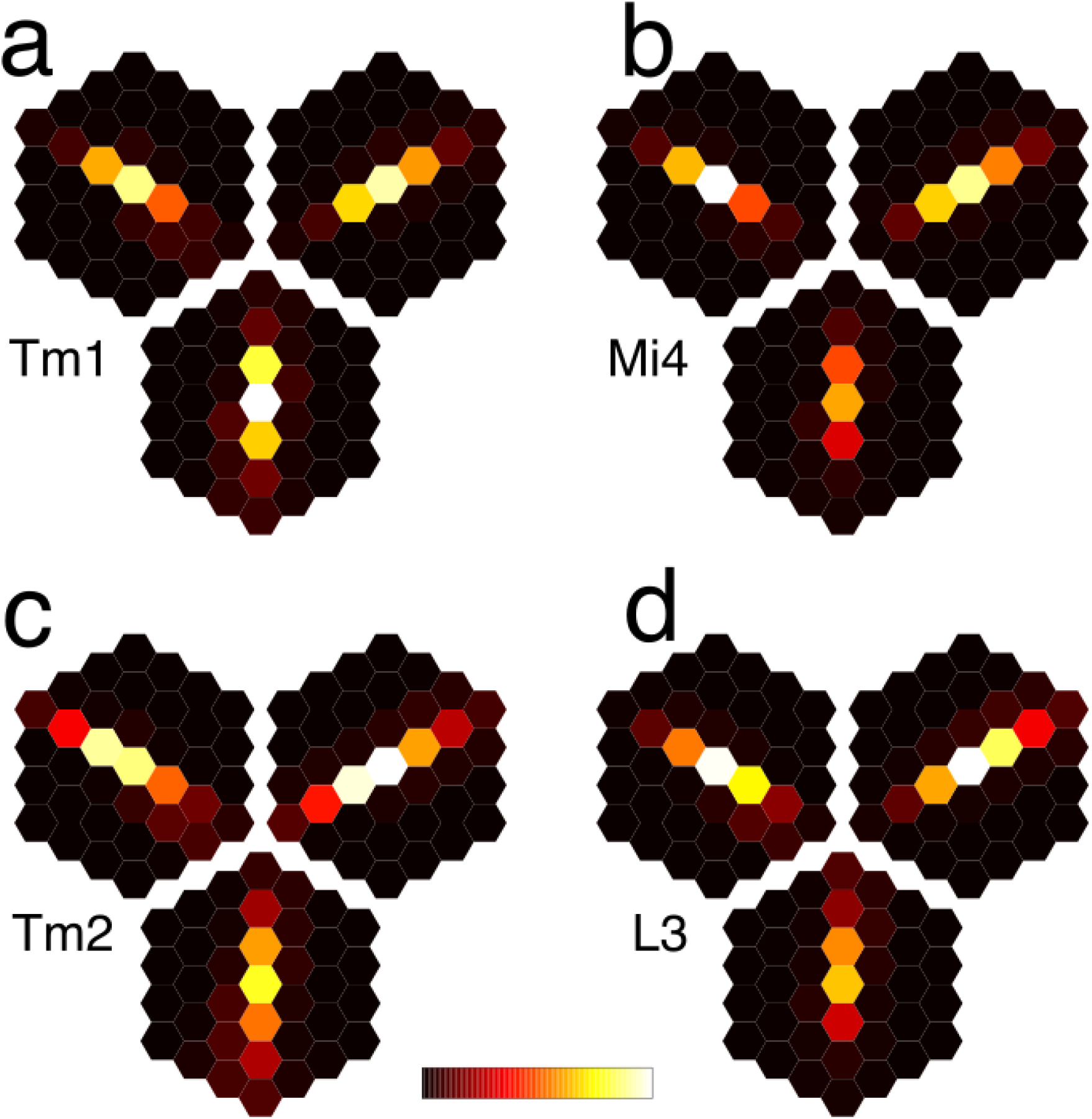
Maps of unicolumnar inputs to Dm3. Each triad of maps is for Dm3v (bottom), Dm3p (top left), and Dm3q (top right). (a) Tm1-Dm3 (b) Mi4-Dm3 (c) Tm2-Dm3 (d) L3-Dm3 Maximum synapse numbers (white in colormap) are 8.2, 4.0, 2.5, and 2.7.

**Figure S4.**
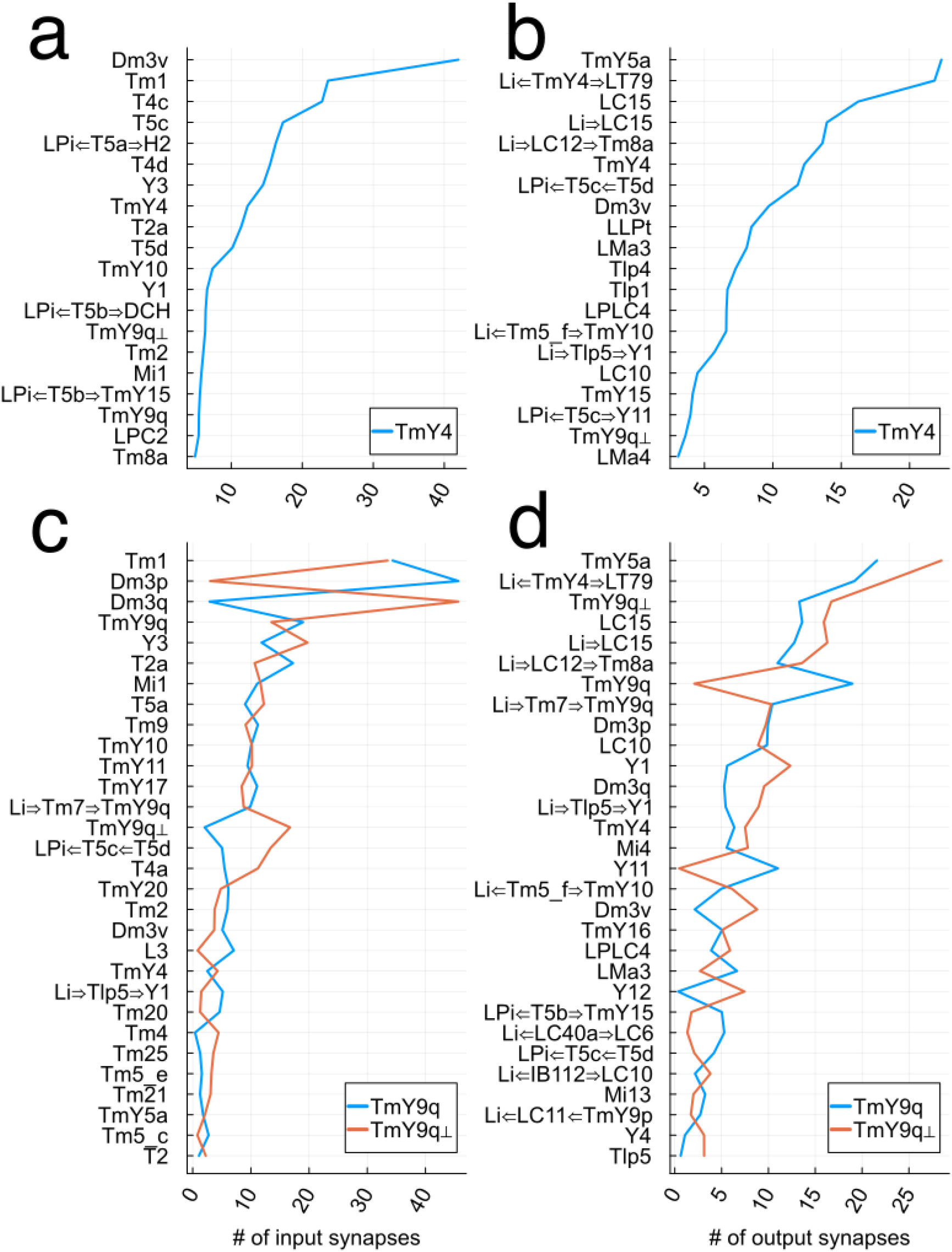
TmY4 and TmY9 inputs and outputs. (a) Average number of input synapses received by a TmY4 cell from presynaptic cell types. (b) Average number of output synapses sent by a TmY4 cell to postsynaptic cell types. (c) Average number of input synapses received by a TmY9 cell from presynaptic cell types. (d) Average number of output synapses sent by a TmY9 cell to postsynaptic cell types.

**Figure S5.**
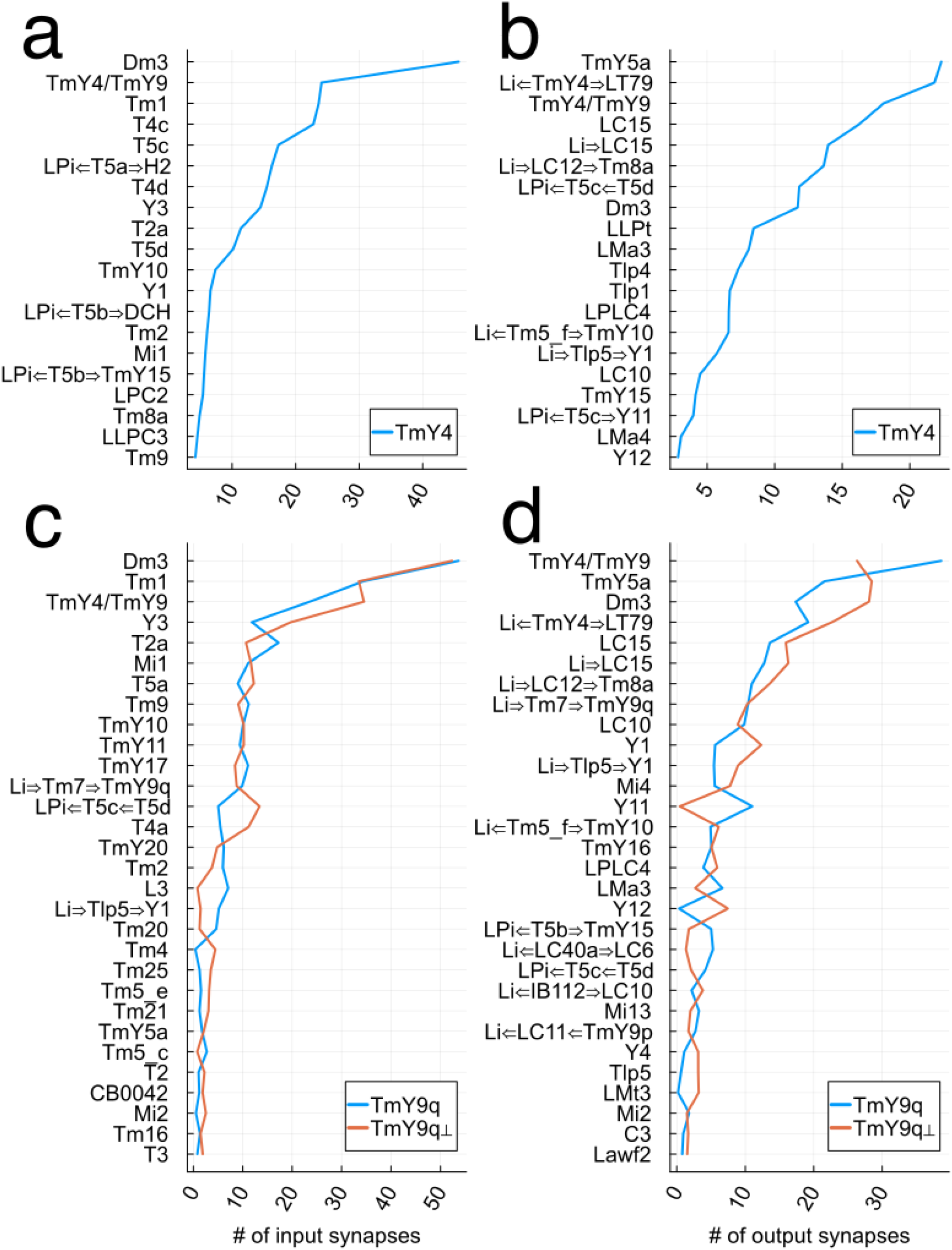
TmY4 and TmY9 inputs and outputs. Same as Figure S4, except Dm3v, Dm3p, and Dm3q are grouped into the presynaptic class Dm3, and TmY4, TmY9q, and TmY9q are grouped into the presynaptic class TmY4/TmY9.

**Figure S6.**
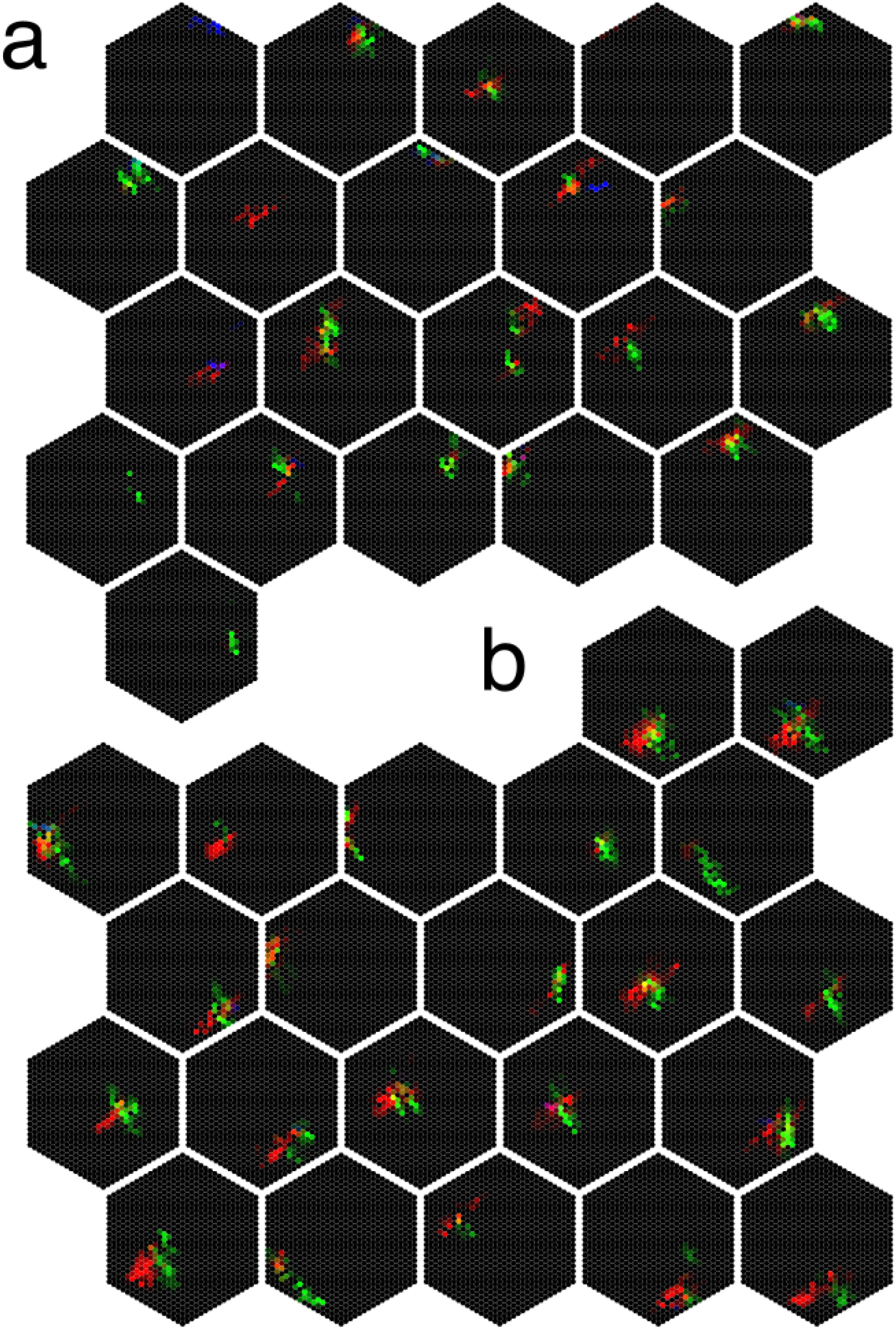
Tm1-TmY-LC10e maps. Inputs to every LC10e cell from Tm1 cells, mediated by TmY9q (red), TmY9q^┴^ (green), and TmY4 (blue) cells. The examples shown in Fig. 5b were drawn from this set of maps for all LC10e cells. The red value for each Tm1 cell is the product of the Tm1-TmY9q and TmY9q-LC10e synapse numbers summed over all intermediate TmY9q cells. Green and blue values are computed analogously. (a) For dorsal LC10e cells, the maps look more noisy. (b) For ventral LC10e cells, the maps often resemble two edges in a corner configuration.

**Figure S7.**
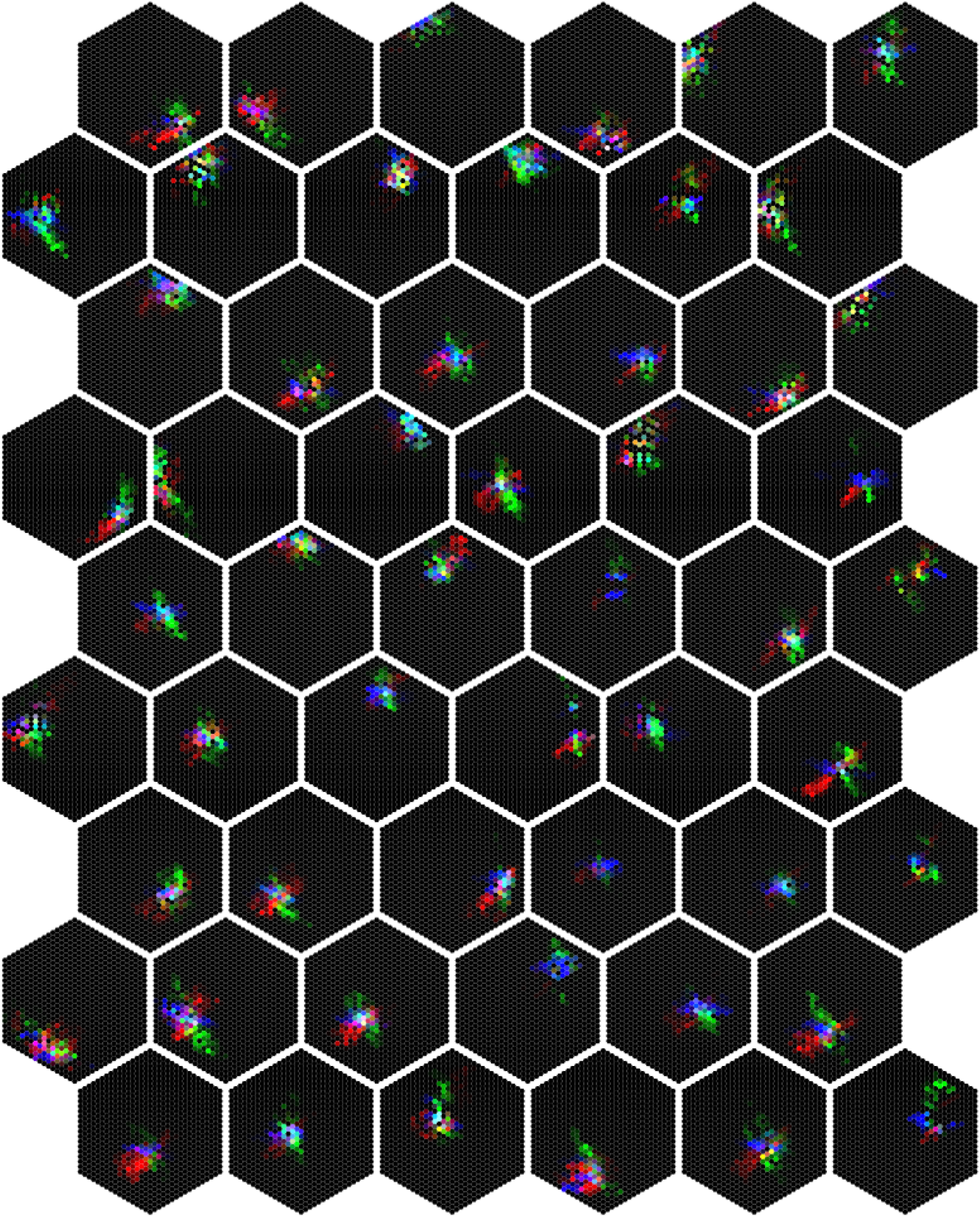
Tm1-TmY-LC15 maps. Inputs to every LC15 cell from Tm1 cells, mediated by TmY9q (red), TmY9q^┴^ (green), and TmY4 (blue) cells. The examples shown in Fig. 5d were drawn from this set of maps for all LC15 cells.

**Figure S8.**
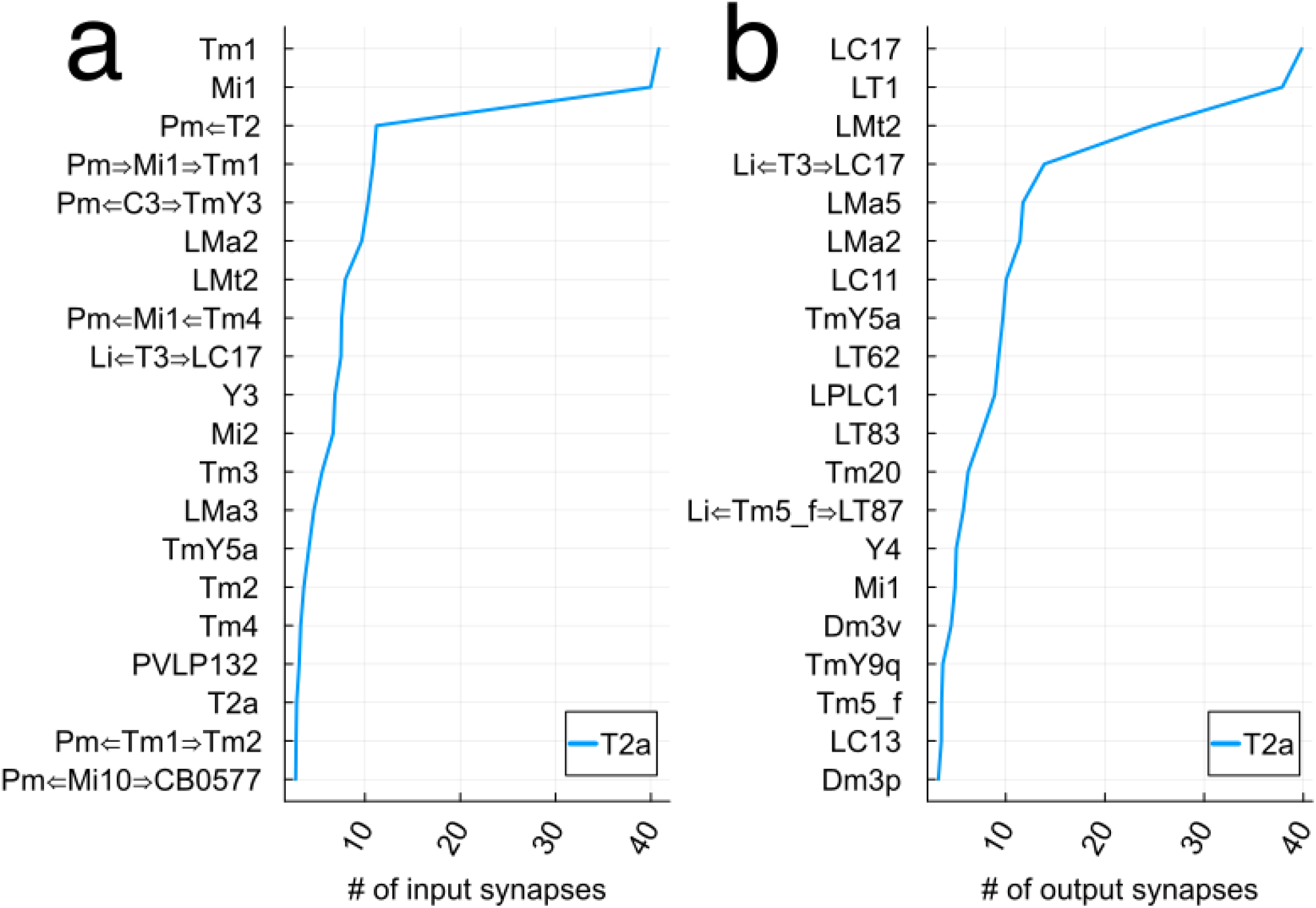
T2a inputs and outputs. (a) Average number of input synapses received by a T2a cell from presynaptic cell types. (b) Average number of output synapses sent by a T2a cell from presynaptic cell types.

**Figure S9.**
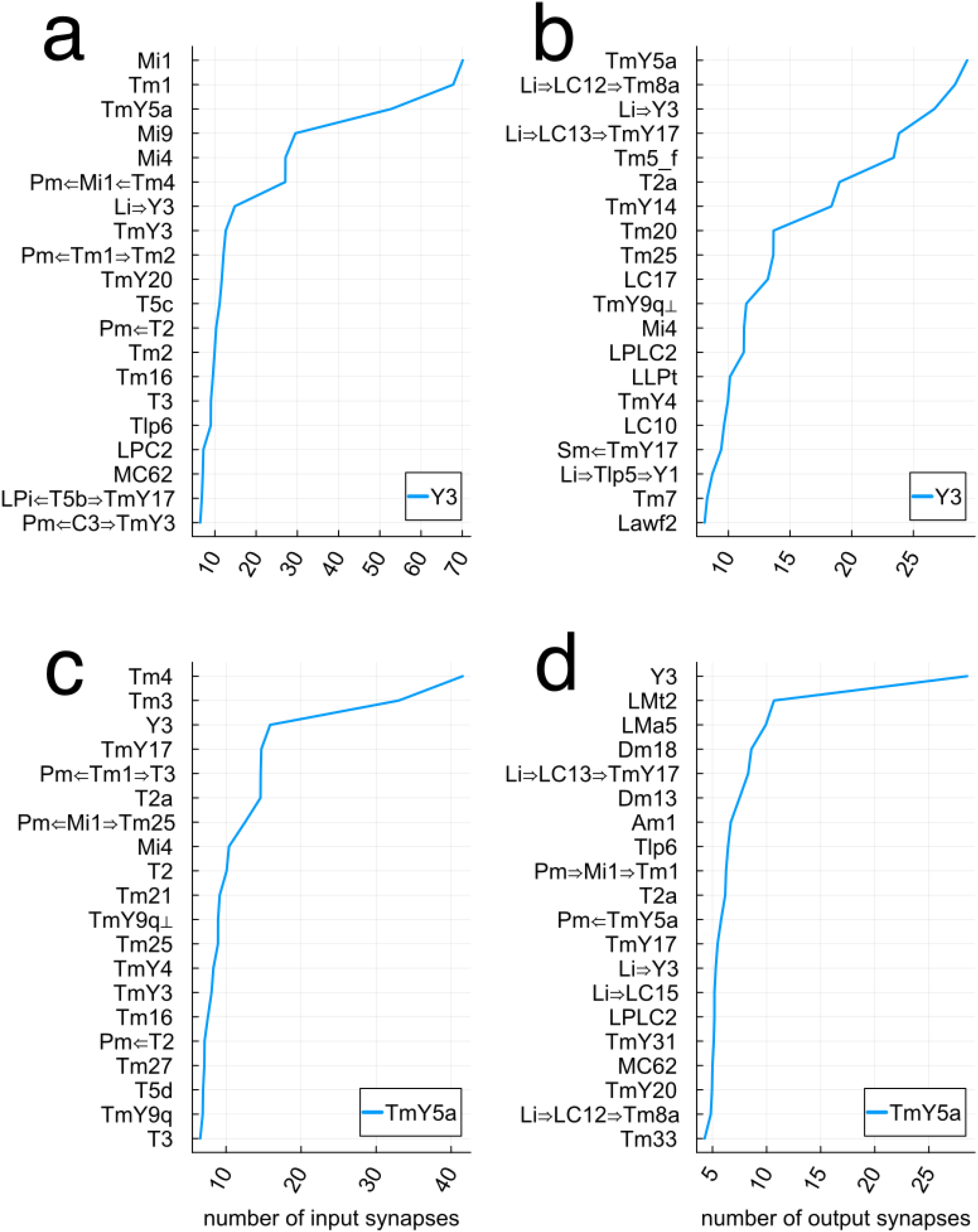
Y3 and TmY5a inputs and outputs. (a) Average number of input synapses received by a Y3 cell from presynaptic cell types. (b) Average number of output synapses sent by a Y3 cell to postsynaptic cell types. (c) Average number of input synapses received by a TmY5a cell from presynaptic cell types. (d) Average number of output synapses sent by a TmY5a cell to postsynaptic cell types.

**Figure S10.**
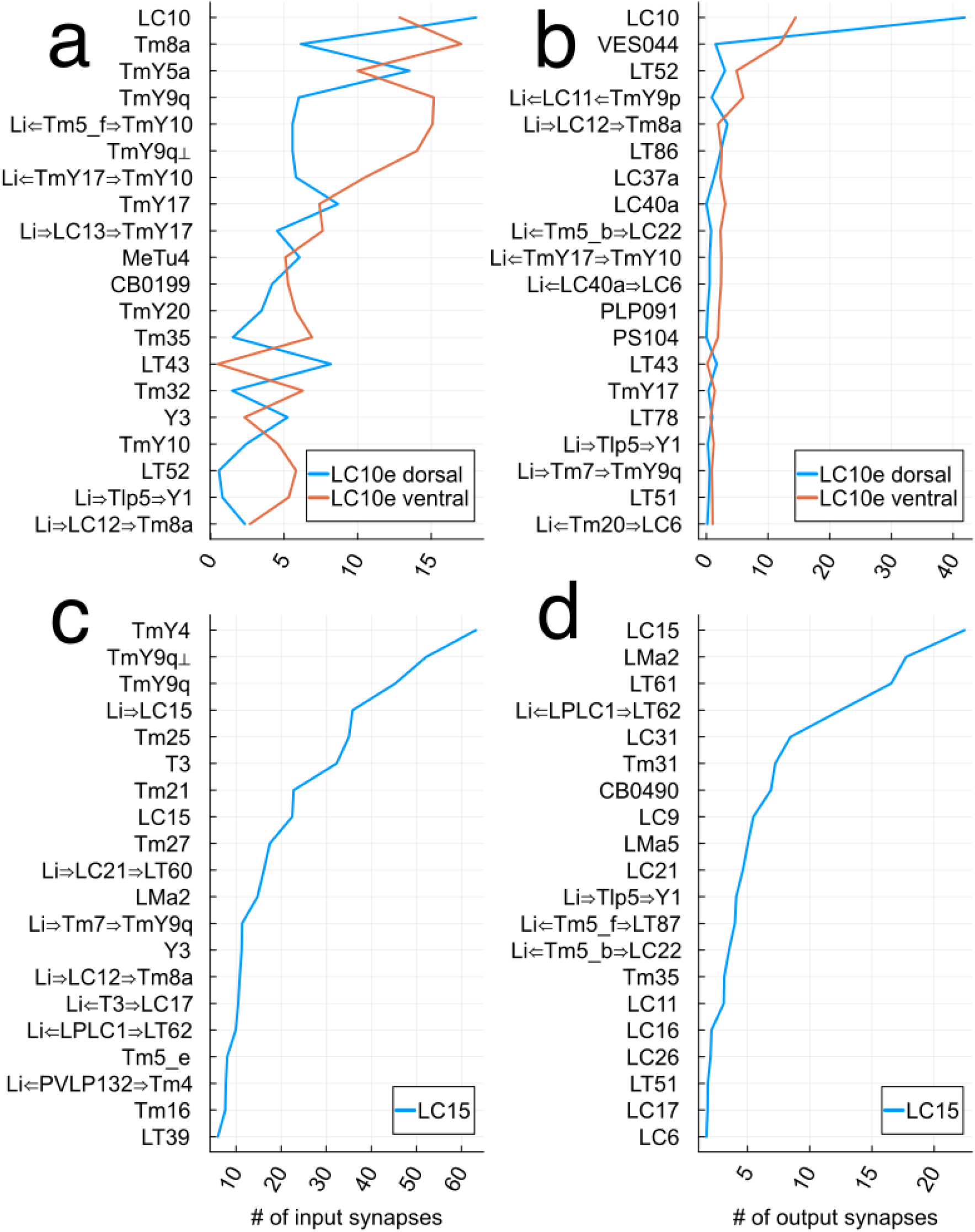
LC10e and LC15 inputs and outputs. Postsynaptic cells are restricted to optic lobe neurons in the graphs, which do not include central brain neurons. (a) Average number of input synapses received by an LC10e cell from presynaptic cell types. (b) Average number of output synapses sent by an LC10e cell to postsynaptic cell types. (c) Average number of input synapses received by a LC15 cell from presynaptic cell types. (d) Average number of output synapses sent by a LC15 cell to postsynaptic cell types.

## Methods

### Spatial coordinates in the optic lobe

A wiring diagram of an adult female *Drosophila* brain is now available (Zheng et al. 2018; Dorkenwald et al. 2023; Schlegel et al. 2023). Neurons intrinsic to the right optic lobe have been classified into cell types, and the rules of connectivity between types are also available (Matsliah et al. 2023). These rules are simplified: whether two cells are connected depends only on the types of the cells, and not on their locations in space.

For a refined characterization of optic lobe connectivity, it is important to assign spatial coordinates to cells. For this purpose, Mi1 is useful because there is one cell per medulla column, and Mi1 cells appear to be in one-to-one correspondence with the ommatidia. All Mi1 cells were semiautomatically assigned to hexagonal lattice points. Locations of L cells were assigned by placing them in one-to-one correspondence with Mi1 cells using the Hungarian algorithm applied to the connectivity matrix. Tm1, Mi4, and Tm2 locations were assigned by placing them in one-to-one correspondence with L cells.

The resulting lattice captures the nearest neighbor relations of the cells and columns, and is not intended to accurately represent distances. A similar lattice can be constructed for ommatidia, and this lattice is left-right inverted relative to the lattice of medulla columns due to the optic chiasm. Therefore back-to-front motion on the retina is front-to-back motion on the medulla lattice. Representations of the lattices that are more spatially accurate can be found in (Zhao et al. 2022).

### Centers of Tm1 input maps

For locations of Dm3, TmY, T2a, and Y3, each Tm1 input map was convolved with a linear filter that was 1.1 in the central column and 1 in its six neighboring columns. The maximum of the result was taken as the center of the Tm1 input map.

## Acknowledgements

I am grateful to Krzysztof Kruk for typing virtually all of the cells used in this analysis (Matsliah et al. 2023), and to Emil Kind for sharing his annotations of Mi1 locations in the right optic lobe. Ari, Nuri, and Seri Seung helped assign Mi1 cells to hexagonal lattice points. I thank Yerbol Kurmangaliyev and Mala Murthy for their comments on the manuscript.

## Notes

https://codex.flywire.ai/app/optic_lobe_catalog

### Competing Interest Statement

The author has a financial stake in Zetta AI.

### Summary of Updates

Each figure caption is now next to the corresponding figure.

